# A 96-well culture platform enables longitudinal analyses of engineered human skeletal muscle microtissue strength

**DOI:** 10.1101/562819

**Authors:** Mohammad E. Afshar, Haben Y. Abraha, Mohsen A. Bakooshli, Sadegh Davoudi, Nimalan Thavandiran, Kayee Tung, Henry Ahn, Howard Ginsberg, Peter W. Zandstra, Penney M. Gilbert

## Abstract

Three-dimensional (3D) in vitro models of human skeletal muscle mimic aspects of native tissue structure and function, thereby providing a promising system for disease modeling, drug discovery or pre-clinical validation, and toxicity testing. Widespread adoption of this research approach is hindered by the lack of an easy-to-use platform that is simple to fabricate and yields arrays of human skeletal muscle micro-tissues (hMMTs) in culture with reproducible physiological responses that can be assayed non-invasively. Here, we describe a design and methods to generate a reusable mold to fabricate a 96-well platform, referred to as MyoTACTIC, that enables bulk production of 3D hMMTs. All 96-wells and all well features are cast in a single step from the reusable mold. Non-invasive calcium transient and contractile force measurements are performed on hMMTs directly in MyoTACTIC, and unbiased force analysis occurs by a custom automated algorithm, allowing for longitudinal studies of function. Characterizations of MyoTACTIC and resulting hMMTs confirms the reproducibility of device fabrication and biological responses. We show that hMMT contractile force mirrors expected responses to compounds shown by others to decrease (dexamethasone, cerivistatin) or increase (IGF-1) skeletal muscle strength. Since MyoTACTIC supports hMMT long-term culture, we evaluated direct influences of pancreatic cancer chemotherapeutics agents on contraction competent human skeletal muscle fibers. A single application of a clinically relevant dose of Irinotecan decreased hMMT contractile force generation, while clear effects on fiber atrophy were observed histologically only at a higher dose. This suggests an off-target effect that may contribute to cancer associated muscle wasting, and highlights the value of the MyoTACTIC platform to non-invasively predict modulators of human skeletal muscle function.

## INTRODUCTION

Skeletal muscle is the most abundant tissue in human body enabling critical physiological and functional activities, such as thermogenesis (Periasamy et al., 2017) and mobility (Lauretani et al., 2003). There are many degenerative and fatal diseases of skeletal muscle that remain untreated and the underlying pathology of some muscle related diseases is not fully understood. The use of animal models to study skeletal muscle diseases has improved our understanding of in vivo drug response and disease pathology (Maltzahn et al., 2012; Thomason and Booth, 1990). However, in some cases animal models fail to accurately predict drug response in humans, in part due to species specific differences leading to inaccurate disease symptoms (Gaschen et al., 1992; McGreevy et al., 2015). Furthermore, animal models are expensive and time consuming making them less desirable for drug testing (DiMasi et al., 2003). As a result, a push to establish in vitro models of human skeletal muscle with reliable phenotypic readouts for drug testing is underway with the goal of improving therapeutic outcomes in humans.

Two-dimensional (2D) cultures of human skeletal muscle cells are most often implemented for drug testing and disease modeling. Despite their ease of use and demonstrated predictive power in certain cases (Stevenson et al., 2005), 2D models of skeletal muscle are ill-suited to in vitro studies of contractile muscle fibers (Bakooshli et al., 2018) by failing to maintain structural integrity over long periods of time (Blau and Webster, 1981; Eberli et al., 2009), and yielding randomly oriented muscle fibers which limits their application for measurement of myofiber contractile force (Smith et al., 2014). A recent report combined 2D culture substrate micropatterning to more closely mimic of the physiological environment, increase reproducibility, and provide an indirect method to assess the contractile capacity of myotubes (Young et al., 2018). This advancement enables scalability and high throughput drug discovery predictions. However, the method is limited in its capacity to maintain muscle fibers long-term, reflected in screens designed to evaluate drugs effects on the earliest phase of differentiation, and analysis is an end-point, thereby precluding longitudinal studies.

Three-dimensional (3D) tissue engineering methods to study skeletal muscle in a dish serve to address these 2D culture gaps, and are beginning to replace conventional assay platforms (Vandenburgh et al., 2008a, 2009). 3D culture models provide multi-dimensional cell-matrix interactions, which is critical to the pathology of conditions such as muscular dystrophies and age-induced muscle fibrosis (Alnaqeeb et al., 1984; Duance et al., 1980). In addition, engineered 3D skeletal muscle models mimic native muscle architecture (Juhas et al., 2014), provide structural integrity for long-term culture of myofibers in vitro, and enable contractile force measurements (Lee and Vandenburgh, 2013). Recent articles report successful development of 3D culture models of human skeletal muscle (Bakooshli et al., 2018; Chal et al., 2016; Madden et al., 2015; Maffioletti et al., 2018; Osaki et al., 2018; Rao et al., 2018). In these studies, active force is quantified on tissues removed from the supporting culture device to implement a force transducer, which is precise, but invasive.

As a non-invasive alternative, others have engineered culture devices that employ flexible posts or pillars to support tissue maturation and measure active force as it relates to post-deflection following electrical or chemical stimulation (Osaki et al., 2018; Sakar et al., 2012; Vandenburgh et al., 2008, 2009). However, some of these technologies have footprints that are not amenable to high-throughput tests. In other cases, device fabrication and/or implementation is perceived as challenging, or make use of cumbersome inserts with vertical posts, all limiting their widespread adoption for drug testing and disease modeling by industry and other researchers.

Here, we describe a method for simple fabrication of a human skeletal muscle (Myo) microTissue Array deviCe To Investigate forCe (MyoTACTIC); a 96-well plate platform for bulk production of human muscle microtissues (hMMTs) for phenotypic drug testing. Our fabrication methodology leads to the highly reproducible single-step casting of a 96 well plate that offers easy workflow integration and requires a relatively small number of cells for tissue formation. We provide an optimized method for formation of functional hMMTs using primary human myogenic progenitor cells in MyoTACTIC by building upon previously published protocols (Bakooshli et al., 2018; Madden et al., 2015). We show that hMMTs self-organize in MyoTACTIC to form multi-nucleated, striated muscle fibers that are responsive to electrical and biochemical stimuli with kinetics and maturation levels matching that observed in larger formats (Madden et al., 2015), and that the process is reproducible from well-to-well of the device.

MyoTACTIC enables easy and non-invasive contractile force and calcium transient measurements of hMMTs over time within the culture device. We demonstrate that known myotoxic drugs (dexamethasone, cerivastatin) induce muscle fiber atrophy and decrease hMMT contractile force generation similar to clinical outcomes, while treating hMMTs with insulin-like growth factor 1 (IGF-1) improves contractile force generation. We then show that a single clinically relevant dose of Irinotecan, a chemotherapeutic reagent used to treat pancreatic cancer, induces muscle fiber atrophy and diminishes contractile force, thereby validating the ability of MyoTACTIC to predict an off-target drug response on human skeletal muscle. We focused our studies on direct muscle fiber effects by initiating all treatments at a time-point when the muscle fibers are fully formed, an assessment made possible by the MyoTACTIC system. Notably, we uncover modified force responses at lower treatment doses than those required to see visible effects by classic histological methods, highlighting the sensitivity of functional metrics over morphological assessments.

In sum, MyoTACTIC enables longitudinal analyses of human skeletal muscle microtissue strength to support fundamental research, drug discovery, and toxicity studies.

## RESULTS

### MyoTACTIC fabrication and implementation is simple and supports bulk production of human skeletal muscle micro-tissues

We report an in vitro platform, hereafter referred to as MyoTACTIC, that supports simple and reproducible culture of contractile human skeletal muscle microtissues (hMMTs) for drug and biomolecule testing. MyoTACTIC is a custom-designed elastomeric 96-well plate (**Figure 1a-c** and **Supplementary Figure 1**) in which each well consists of an elliptical inner chamber containing two posts at either end (**Figure 1c,e-g** and **Supplementary Figure 1**). A multi-step casting process is employed to fabricate MyoTACTIC plates (**Figure 1a**) from a 3D printed design (see Materials and Methods). The fabrication process leads to the generation of a reusable polyurethane (PU) negative mold for reproducible generation of MyoTACTIC plates containing 96-wells and all well features using single step polydimethylsiloxane (PDMS) casting within 3 hours (**Figure 1a**, Step 4).

**Figure 1.**
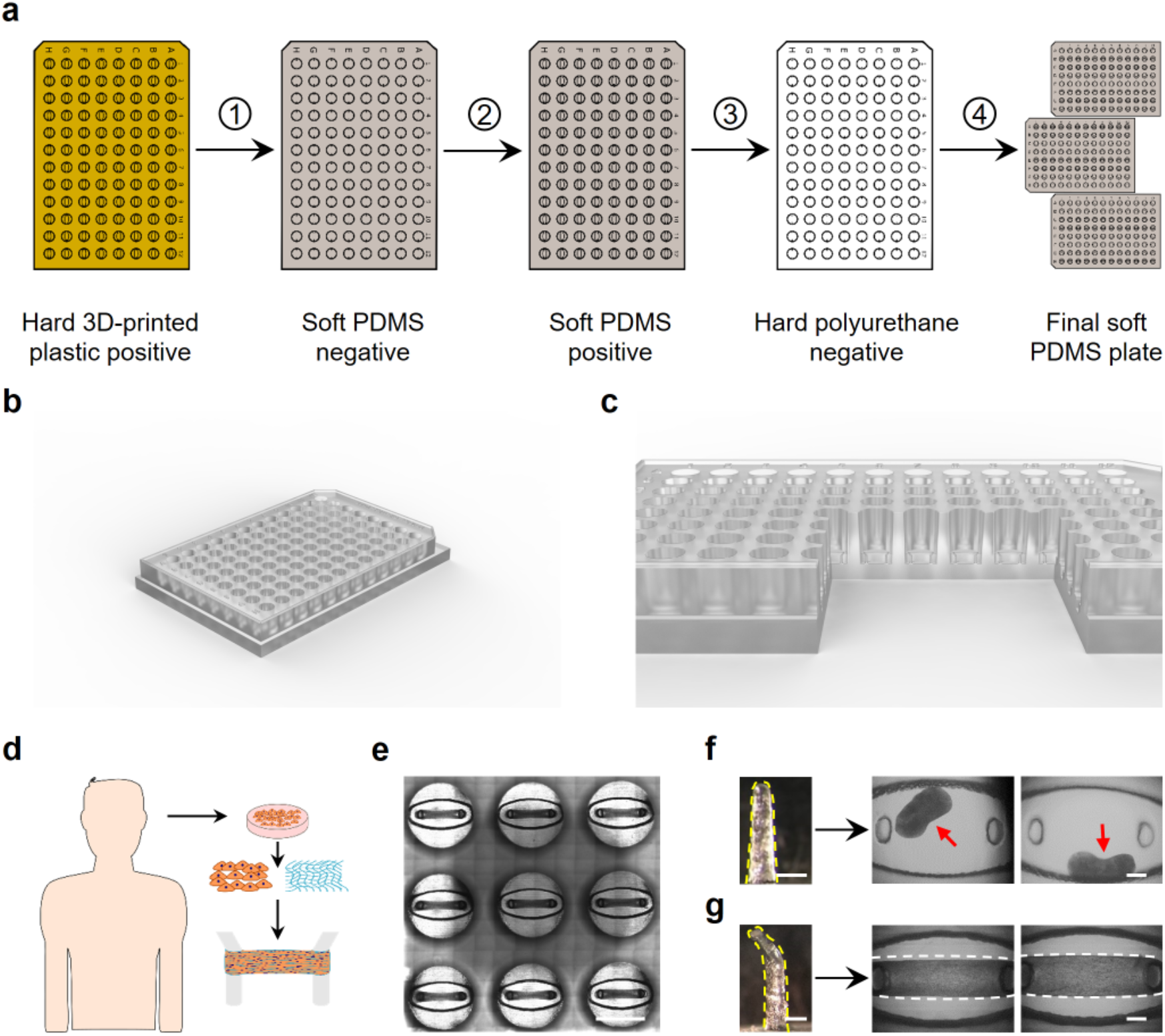
Design and production of the MyoTACTIC platform. **(a)** MyoTACTIC production started with creating a three-dimensional Computer Aided Design (3D CAD) using SolidWorks™ Software which was then printed using a ProJet MJP 3500 3D printer from 3D SYSTEMS. Next, a negative PDMS mold was created from the 3D printed part which was subsequently used to fabricate a soft replica of the design after silanizing. Finally, a rigid negative polyurethane mold was created form the PDMS replica which was used to fabricate multiple PDMS plates. **(b-c)** Computer generated 3D images of MyoTACTIC 96-well plate design and **(c)** a cross-section of wells indicating the location of the micro-posts. **(d)** Schematic overview of human cell isolation and subsequent generation of hMMTs in MyoTACTIC. **(e)** Stitched phase-contrast image of 9 wells of MyoTACTIC containing remodeled hMMTs 10 days post seeding. Scale bar 5 mm. **(f-g)** Impact of micro-post design on formation and long-term maintenance of hMMTs in MyoTACTIC. Representative images of **(f)** collapsed and **(g)** successfully remodeled hMMTs seeded in wells with **(f)** hook-less and **(g)** hook featured posts. Micro-posts are outlined in yellow dashed lines. Red arrows indicate collapsed hMMTs on the top right panels and hMMTs are outlined in white dashed lines on the bottom right panels. Scale bars 500 μm.

Three-dimensional (3D) hMMTs are engineered using purified primary human myogenic progenitor cells (**Figure 1d** and **Supplementary Figure 2a**) suspended in a hydrogel mix (**Table S2**) based on previously published work (Bakooshli et al., 2018; Madden et al., 2015), by pipetting the cell-hydrogel suspension into the MyoTACTIC well chambers, between and around the posts (**Figure 1c-e** and **Figure 2a, left panel**). The micro-posts included in each well serve as tendon-like anchor points to establish uniaxial tension in the remodeling hMMT (**Figure 1c,e-g**) and direct micro-tissue formation and compaction in each well (**Figure 1c,e,g** and **Supplemental Figure 2b**).The hook feature at the top each post is an essential design criteria as hMMTs migrate off the hook-less posts within two days of cell seeding (**Figure 1f**). Indeed, MyoTACTIC design (e.g. post size, shape, and positioning, cell seeding chamber size and shape, platform material stiffness, etc.) was iterated to enable bulk production of hMMT arrays amenable to the ‘in dish’ functional analyses described herein.

**Figure 2.**
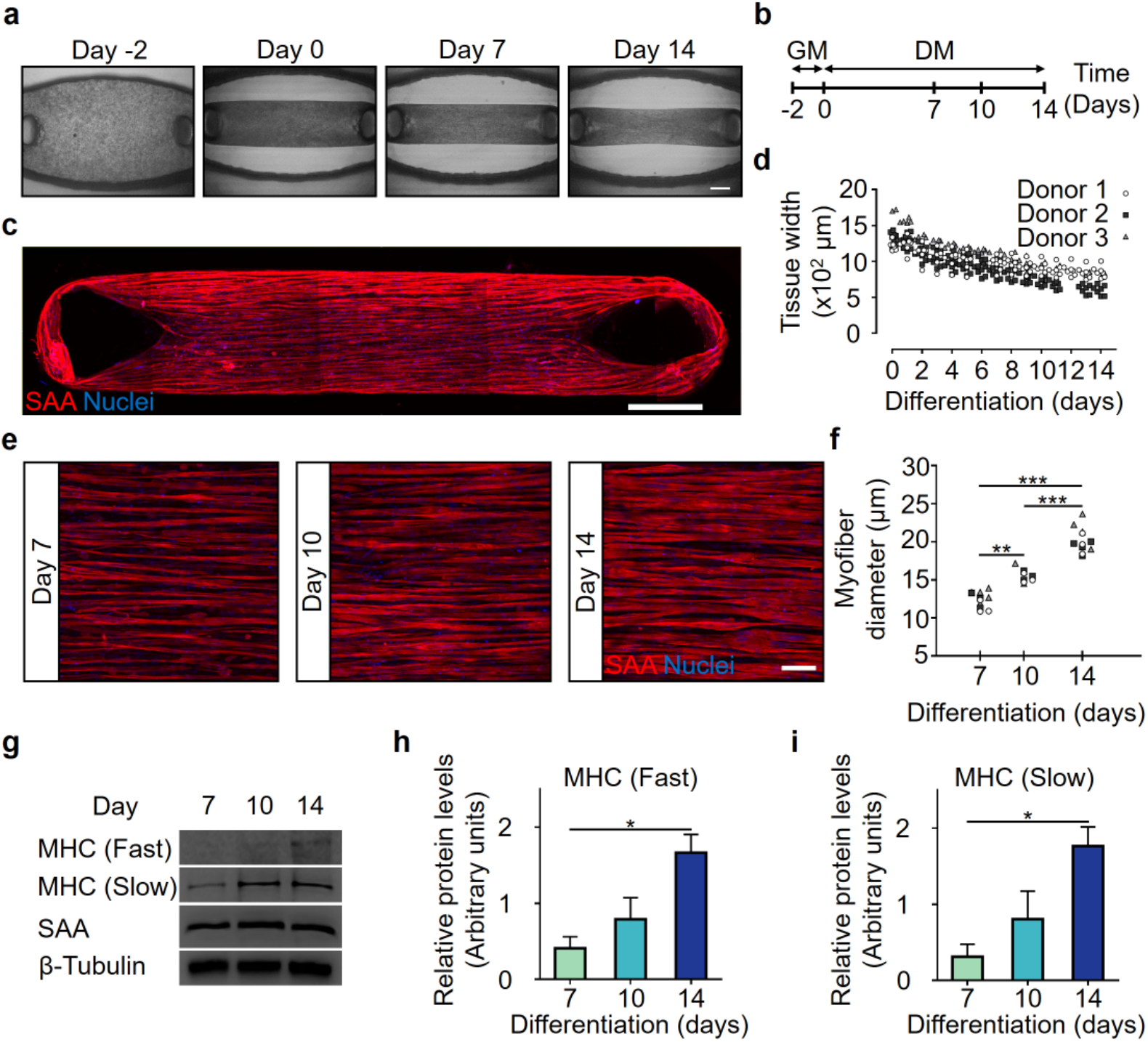
MyoTACTIC enables formation of hMMTs with aligned myofibers exhibiting hypertrophy and adult myosin heavy chain expression. **(a)** Representative phase-contrast images of hMMTs depicting the remodeling of the extracellular matrix by human myoblasts over time. Day 0 marks the time for switching to differentiation media. Scale bar 500 μm. **(b)** Schematic diagram of the timeline for hMMT culture. hMMTs are cultured in growth media (GM) for the first two days (day −2 to day 0) and then the media is switched to differentiation media (DM) on Day 0. **(c)** Representative confocal stitched image of a hMMT cultured for 2 weeks, immunostained for sarcomeric α-actinin (SAA, red) and exposed to DRAQ5 (blue) to counter stain the nuclei. Scale bar 500 μm. **(d)** Dot plot indicating the width of hMMTs over the course of culture time. (n = minimum of 16 hMMTs from 3 muscle patient donors per time point). **(e)** Representative confocal images of muscle fibers formed in hMMTs and immunostained for SAA (red) and nuclei (blue) after 7, 10, and 14 days in differentiation media. Scale bar 50 μm. (f) Quantification of hMMT fiber diameter over time. **p < 0.01, *** p < 0.001 (n = minimum of 9 hMMTs from 3 muscle patient donors per time point). **(g)** Representative western blot images of myosin heavy chain (MHC) isoforms (fast and slow), SAA, and β-tubulin over culture time (day 7 to 14) and **(h-i)** bar graph quantification of relative **(h)** MHC-fast and **(i)** MHC-slow protein expression in hMMTs over time. *p < 0.05 (n = 3 blots from 3 muscle patient donors per time point, where each blot is run using lysate of 4 hMMTs from a single patient donor lysed together). In **(d)** and **(f)** each symbol represents data from one patient donor. In **(h)** and **(i)** values are reported as mean ± SEM. In **(f)**, **(h)**, and **(i)** significance was determined by one-way ANOVA followed by multiple comparisons to compare differences between groups using Tukey’s multiple comparisons test.

### Human muscle microtissues cultured in MyoTACTIC exhibit structural characteristics similar to macro-tissues

In order to ensure that hMMTs cultured in MyoTACTIC possess structural characteristics comparable to previously published methods (Bakooshli et al., 2018; Madden et al., 2015), in spite of using fewer cells and a miniaturized format, we investigated the structural characteristics of hMMTs in culture over time. hMMTs remodeled within two days of culture in myogenic growth media without bFGF (**Figure 2a-b** and **Table S2**) and continued to remodel and compact over the additional two-weeks in differentiation media (**Figure 2a-b,d** and **Supplementary Figure 3a**).

Within two weeks of differentiation, hMMTs formed multi-nucleated and aligned muscle fibers as evident by sarcomeric α-actinin (SAA) immunofluorescence analysis (**Figure 2c**). The vast majority of cells were post-mitotic by Day 7 of differentiation (**Supplementary Figure 3b**), and we also identified formation of cross-striated muscle fibers containing clustered acetylcholine receptors (AChRs) at this time point, which persisted until at least Day 14 (**Supplementary Figure 3c**). We quantified myofiber width at Days 7, 10 and 14 of differentiation (**Figure 2e-f**). As expected, myofiber width increased with time in culture. We noted that hMMT myofiber hypertrophy is sensitive to and supported by autocrine cues, underlying the importance of reserving a portion of the hMMT conditioned media during media exchanges (**Supplementary Figure 3d**). Finally, structural maturation of the hMMTs in culture was evident by significantly higher expression of the adult subtypes (fast and slow) of the contractile protein myosin heavy chain (MHC) at the later stages of culture time (**Figure 2g-i**), while SAA protein expression remained relatively steady over time (**Figure 2g**).

Our analysis demonstrates that hMMTs cultured in MyoTACTIC exhibit similar remodeling dynamics, structural characteristics, and maturation levels to their macroscale counterparts. Further, since some variation is observed between biological replicates, but variation amongst technical replicates is negligible (**Figure 2d,f**), we conclude that the process of hMMT development within MyoTACTIC is reproducible.

### MyoTACTIC allows for non-invasive and in situ hMMT contractile force assessment

hMMTs cultured in MyoTACTIC exhibited spontaneous contractions in culture between Days 10 to 12 of differentiation (**Supplementary Movie S1**), and were competent to produce twitch and tetanus contractions in response to electrical stimuli (0.5 Hz and 20 Hz respectively; **Supplementary Movie S2** and **Supplementary Figure 4**). As predicted based on the presence of AChR clusters (**Supplementary Figure 3c**), hMMTs generated an immediate and robust tetanus contraction in response to biochemical (ACh, 2mM) stimulation (**Supplementary Movie S3**). These observations confirm hMMT functional maturation in MyoTACTIC.

In vitro models of human skeletal muscle are generally incapable of studying muscle contraction, and those that can are limited to doing so as an experimental endpoint owing to the need to remove tissues from the culture device so as to implement a force transducer for measurements (Madden et al., 2015; Young et al., 2018). MyoTACTIC micro-posts were designed to sustain hMMT long-term culture and to allow for non-invasive contractile force measurements of hMMTs in a fast and reproducible manner (**Figure 3a**). Mechanical analysis of MyoTACTIC micro-posts confirmed a linear force-displacement relation (**Figure 3a-b**) in saline (37 °C). This material property allows for hMMT contractile force to be determined by comparing the extent of post deflection (Vandenburgh et al., 2008) induced by hMMTs during contraction in response to electrical and biochemical stimuli (**Figure 3a-b**).

**Figure 3.**
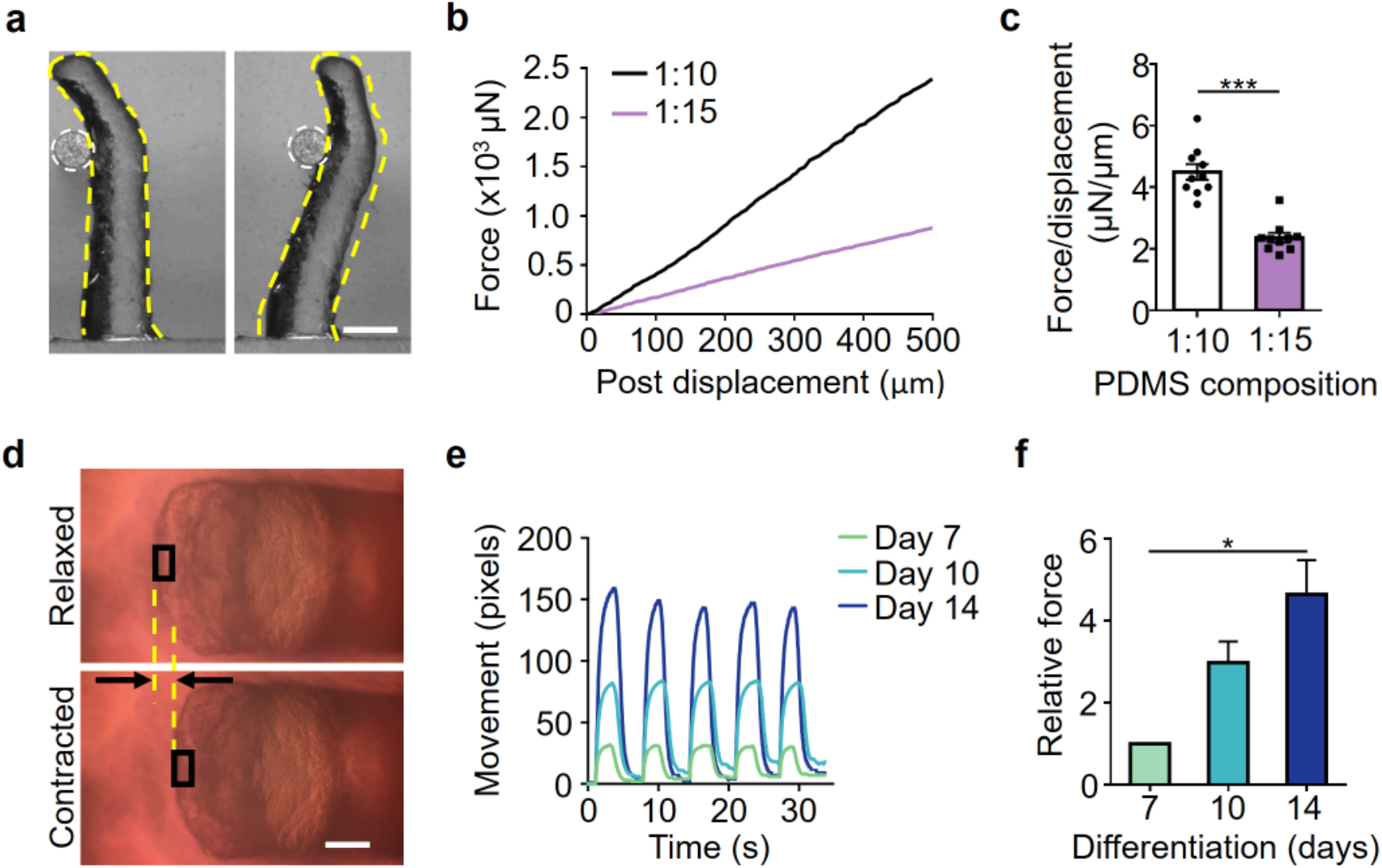
MyoTACTIC enables non-invasive and in situ measurement of hMMT contractile force. **(a)** Phase-contrast images of a micro-post (outlined in yellow dashed lines) displaced in response to the force exerted by a microwire (outlined in white dashed circles). Scale bar 500 μm. **(b)** Plot depicting the relation between force and displacement of micro-posts fabricated using two different PDMS compositions (curing agent: monomer), as measured by the Microsquisher. **(c)** Bar graph quantification of the average force/displacement ratio for the micro-posts formed using two different PDMS compositions. ***p < 0.001 (n = 10 micro-posts per condition). **(d)** Representative bright-field images of a micro-post under 10X magnification before (relaxed) and during (contracted) a tetanus contraction using 20 Hz electrical stimuli. Yellow dashed lines and arrows indicate the displacement of the post. Scale bar 200 μm. **(e)** Representative line graph traces of the micro-post displacement during high frequency (20 Hz) electrical stimulation of hMMTs at Day 7, 10, and 14 of differentiation measured by the custom-written Python computer vision script. **(f)** Bar graph quantification of the relative force generated by hMMTs at Day 7, 10, and 14 of differentiation. Values are normalized to Day 7. *p < 0.05 (n = minimum of 11 hMMTs from 3 muscle patient donors per time point). In **(c)** and **(f)** values are reported as mean ± SEM. Significance was determined by t-test in (c) or Kruskal-Wallis test followed by Dunn’s multiple comparisons test to compare differences between groups in **(f)**.

Since the mechanical modulus of the micro-post design can be modified by adjusting the PDMS monomer to curing agent ratio (**Figure 3c**), it is possible to tune micro-post deflection properties (**Figure 3b**). To maximize post deflection, which in turns serves to minimize measurement error, we implemented the lower curing agent to monomer ratio (1:15) for the force measurements in this study. Notably, well-to well-variation in micro-post mechanical properties was minimal (**Figure 3c**). Next, we developed a post tracking script in Python to non-invasively and in situ quantify hMMT strength through analysis of post deflection videos (**Figure 3d, Supplementary Movie S4**, and **Appendix**). Using the semi-automated and unbiased post tracking script, we confirm that the contractile force of the hMMTs increases significantly with culture time from Day 7 to 14 of differentiation (**Figure 3e-f** and **Supplementary Movie S5**).

Here we conclude that MyoTACTIC, together with our post tracking script, provides a powerful system for longitudinal phenotypic studies of hMMT force generation.

### MyoTACTIC enables non-invasive and in situ hMMT calcium transient assessment

To evaluate the calcium handling properties of hMMTs in MyoTACTIC, we generated microtissues using GCaMP6 stably transduced human muscle progenitor cells (Chen et al., 2013), a sensitive calcium indicator protein, driven by the MHCK7 (Madden et al., 2015) promoter, a muscle specific gene. hMMTs exhibited spontaneous myofiber calcium transients (**Supplementary Movie S6**) in as early as Day 7 of differentiation. GCaMP6 signals arising from hMMTs are captured in the context of MyoTACTIC, allowing for assessment over time. hMMTs generated strong collective calcium transient in response to electrical stimulation (**Figure 4a-c** and **Supplementary Movies S7**) and immediately following exposure to ACh (**Figures 4a,d** and **Supplementary Movie S8**). The magnitude of stimulated calcium transients increased significantly with culture time as measured by normalized fluorescence intensity (ΔF/F_0_; **Figure 4b-e**). Furthermore, pre-treatment with d-tubocurarine (25 μM) blocked ACh-induced calcium transients (**Figure 4f** and **Supplementary Movie S9**), but had no significant effect on hMMT calcium transients induced by electrical stimuli (**Figure 4f** and **Supplementary Movie S9**), which mimics the in vivo muscle response (Ballantyne and Chang, 1997; Bowman, 2006), and validates the functional activity of AChR clusters formed on hMMTs (**Supplemental Figure 3c**).

**Figure 4.**
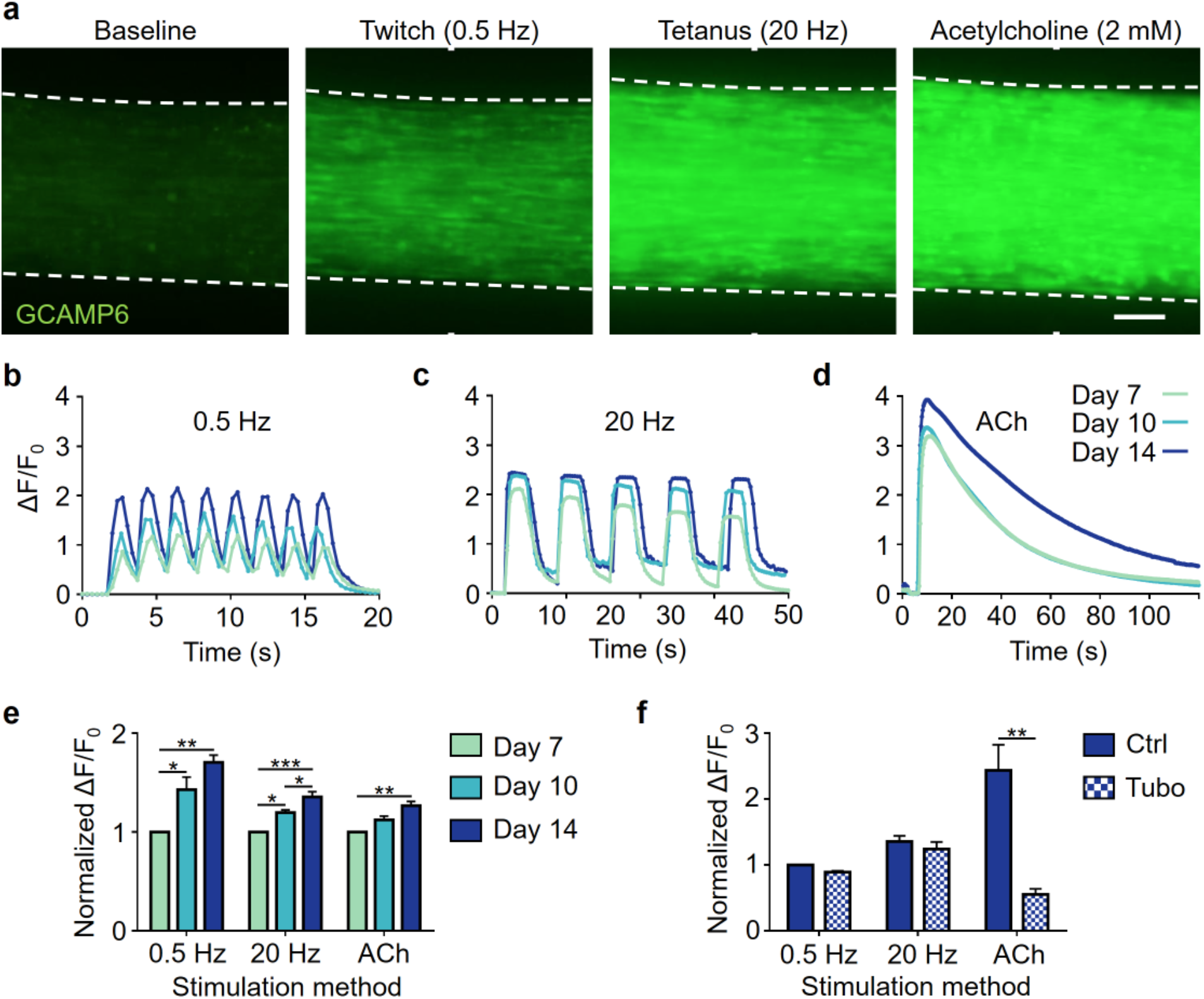
MyoTACTIC enables non-invasive and in situ measurement of hMMT calcium transients. **(a)** Representative epifluorescence images of the peak GCaMP6 signal from hMMTs in response to low frequency (0.5 Hz, twitch contraction), high frequency (20 Hz, tetanus contraction) and acetylcholine (ACh, 2 mM) stimulations at Day 14 of differentiation. Scale bar 200 μm. **(b-d)** Representative calcium transient traces of hMMTs at Day 7, 10, and 14 of differentiation in response to **(b)** low and **(c)** high frequency electrical and **(d)** acetylcholine stimulations. **(e)** Bar graph quantification of hMMTs calcium transients in response to electrical (0.5 and 20 Hz) and biochemical (ACh) stimuli at differentiation Day 7, 10 and 14. Values are normalized to the Day 7 results for each stimulation modality. *p < 0.05; **p < 0.01, *** p < 0.001 (n = minimum of 9 hMMTs from 3 muscle patient donors per time point, per stimulation method). **(f)** Bar graph quantification of calcium transients in hMMTs activated with electrical or biochemical stimuli following pre-treatment with d-tubocurarine (25 μM) at Day 14 of differentiation. Values are normalized to day 14 control hMMTs. **p < 0.01 (n = 9 hMMTs from 3 muscle patient donors per treatment condition per stimulation method). In **(e)** and **(f)** values are reported as mean ± SEM. Significance was determined by two-way ANOVA followed by Tukey’s and Sidak’s multiple comparisons to compare differences between groups in **(e)** or t-test in **(f)**.

### MyoTACTIC cultured hMMTs predict structural and functional treatment responses

Accurate drug response prediction is a crucial requirement if engineered tissues are to be implemented for drug testing. Hence, we studied hMMT myofiber size and contractile strength responses to treatment with three compounds with well-studied effects on skeletal muscle. Since MyoTACTIC supports long-term culture, compounds were administered from Day 7 to 14 of culture to evaluate treatment effects on multinucleated myofiber morphology and function. Immunofluorescence analysis confirmed that dexamethasone and cerivastatin treatments, which are known to induce rhabdomyolysis in humans (Staffa et al., 2002; Tamraz et al., 2013; Viguerie et al., 2012), elicited a dose-dependent decrease in myofiber width (**Figure 5a-b, left** and **middle panels**). In contrast, hMMTs treated with IGF-1 displayed no change in myofiber size, regardless of dose (**Figure 5a-b, right panels**). We then studied compound treatment effects on function by assessing hMMT contractile force. As predicted by morphological assessment, dexamethasone and cerivastatin treatments induced contractile weakness (**Figure 5c, left** and **middle panels** and **Supplementary Movies S10-S11**). Notably, IGF-1 treated (100 nM) hMMTs exhibited significant contractile force gain compared to control (**Figure 5c right panel** and **Supplementary Movie S12**) akin to in vivo animal studies (Coleman et al., 1995; Musarò et al., 2001), and suggesting that MyoTACTIC is capable of perceiving compound effects through non-invasive force analysis that might otherwise be overlooked using more conventional methods (Semsarian et al., 1999).

**Figure 5.**
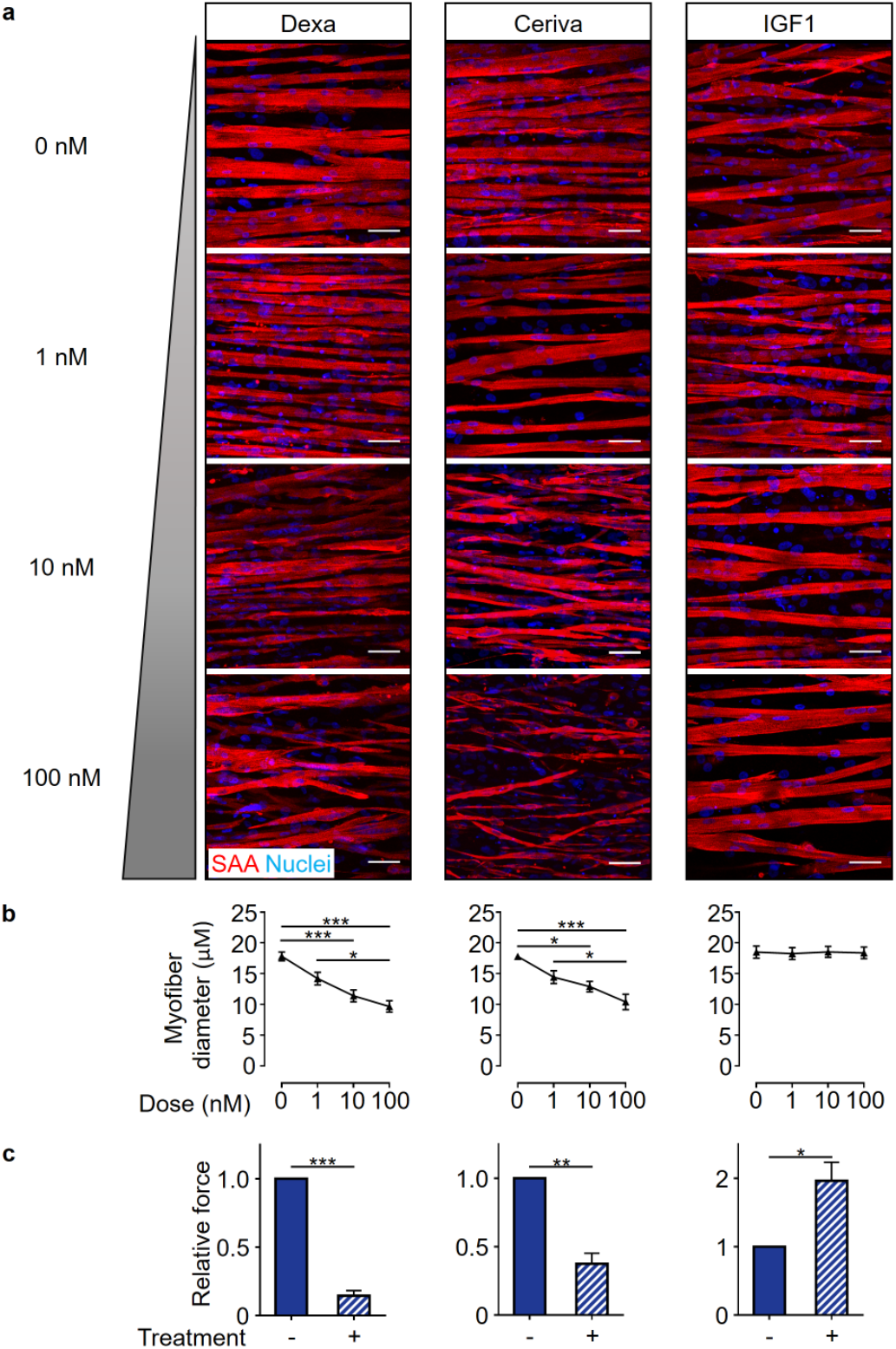
MyoTACTIC-cultured hMMTs predict skeletal muscle structural and functional responses to pharmacological treatment. **(a)** Representative confocal images of Day 14 hMMTs treated for 7 days with either DMEM control or increasing doses (1, 10, and 100 nM) of Dexamethasone (left panels), Cerivastatin (middle panels), or IGF-1 (right panels) and immunostained for sarcomeric a-actinin (SAA, red) and Hoechst 33342 (Nuclei, blue). Scale bar 50 μm. **(b)** Quantification of the dose-dependent effect of Dexamethasone (left panel), Cerivastatin (middle panel), and IGF-1 (right panel) on hMMT myofiber diameter. *p < 0.05, ***p < 0.001 (for each treatment, n = minimum of 4 hMMTs from a minimum of 3 muscle patient donors per treatment dose). **(c)** Bar graph quantification of the tetanus (20 Hz electrical stimuli) contractile force generated by hMMTs treated from Day 7 to 14 with either vehicle (−) or (+) Dexamethasone (10 nM; left panel), Cerivastatin (10 nM; middle panel), and IGF-1 (100 nM; right panel). Ethanol, ddH_2_O, and 10 mM HCl were vehicle controls for Dexamethasone, Cerivastatin, and IGF-1 respectively. Values are normalized to vehicle-treated results. *p < 0.05, ***p < 0.001 (for each treatment, n = minimum of 6 hMMTs generated from 3 muscle patient donors per treatment condition). In **(b)** and **(c)** values are reported as mean ± SEM. In **(b)**, significance was determined by one-way ANOVA followed by Tukey’s multiple comparisons to compare differences between groups (Dexamethasone and Cerivastatin) or Kruskal Wallis test followed by Dunn’s multiple comparisons to compare differences between groups (IGF-1). In **(c)**, significance was determined by t-test (Dexamethasone and Cerivastatin) or t-test with Welch’s correction (IGF-1).

Together, our data demonstrate that hMMT treatments in MyoTACTIC accurately reflect the known effects and can be evaluated via a non-invasive metric of function.

### MyoTACTIC study predicts direct effect of chemotherapeutic drug on human skeletal muscle

Given the predictive response of hMMTs to treatments with known effects on skeletal muscle, we next applied the system to interrogate potential skeletal muscle off-target effects of cancer chemotherapeutic drugs. Cancer-induced skeletal muscle wasting, known as cachexia, has a poorly understood pathology. Cachexia is emerging as an independent determinant of patient survival (Argilés et al., 2014; Dimitriu et al., 2005), suggesting treatment strategies designed with skeletal muscle health in mind are desirable. Since cancer cells are characterized in part by deregulated proliferation, many commonly used chemotherapeutic agents non-selectively target cycling cells. Skeletal muscle fibers are post-mitotic, hence, the possibility of a direct effect of chemotherapeutic agents on muscle health is underexplored. Patients with pancreatic cancer are strongly associated with cachexia (Argilés et al., 2014; Dodson et al., 2011; Sun et al., 2015), therefore we selected Gemcitabine and Irinotecan for MyoTACTIC analysis, both of which employ mitosis-targeting mechanisms of action (Keil et al., 2015; Moysan et al., 2013), and are used to treat patients with advanced pancreatic cancer.

Drug doses were selected based on reported clinical concentrations (Chabot, 1997; Conroy et al., 2011), in addition to evaluating a supraphysiological dose for each drug. The treatment began on Day 7 of differentiation, when multinucleated myofibers are prevalent (**Figure 2**) and functional (**Figures 3–4**), and few mitotic cells remain (**Supplementary Figure 3b**). The regimen in this study was a single dose to model a typical clinical treatment, followed by analysis one week later. Gemcitabine, regardless of dose (32 μM and 320 μM), had no significant effect on hMMT myofiber structural organization (**Figure 6a**) or contractile force generation (**Figure 6b** and **Supplementary Movie S13**). Conversely, Irinotecan exposure elicited a dramatic reduction in hMMT strength at the clinical dose (16 μM) with clear effects on myofiber integrity only visible histologically at the supraphysiological dose (**Figure 6c-d** and **Supplementary Movie S14**).

**Figure 6.**
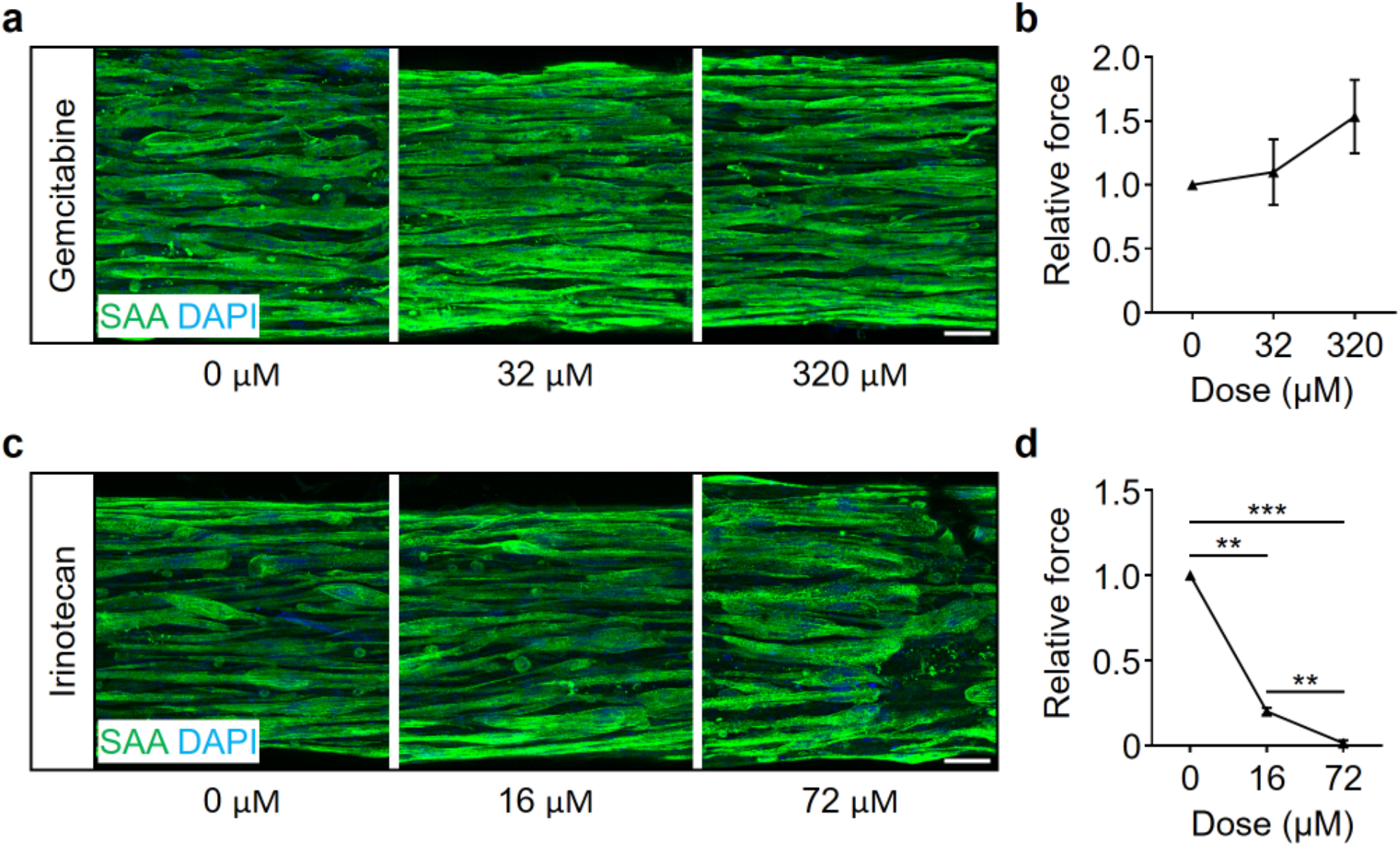
MyoTACTIC cultured hMMTs predict direct effect of chemotherapeutic agents on human skeletal muscle contractile function. **(a, c)** Representative confocal images of Day 14 hMMTs treated with one-time administration of vehicle (DMSO for Irinotecan and PBS for Gemcitabine) or increasing doses of **(a)** Gemcitabine or **(c)** Irinotecan on Day 7 and then immunostained for sarcomeric α-actinin on Day 14. Scale bar 50 μm. **(b, d)** Quantification of relative tetanus contractile force generation by Day 14 hMMTs treated with vehicle (DMSO for Irinotecan and PBS for Gemcitabine), **(b)** Gemcitabine (32 μM and 320 μM), or **(d)** Irinotecan (16 μM and 72 μM). Values in **(b)** and **(d)** are normalized to vehicle treated results. **p < 0.01, ***p < 0.001 (n = minimum of 5 hMMTs from 3 biological replicates sourced from 2 human muscle patient donors per treatment condition). In **(b)** and **(d)** values are reported as mean ± SEM. Significance was determined by one-way ANOVA followed by Tukey’s multiple comparisons to compare differences between groups in **(b)** and **(d)**.

This study extolls the capacity of MyoTACTIC to predict unexpected off-target effects on skeletal muscle force generation with a greater sensitivity than standard histological methods.

## DISCUSSION

Here we report a design and methods to fabricate a 96-well culture platform, MyoTACTIC, for the bulk production of hMMTs, and we show the feasibility of hMMT implementation in phenotypic compound testing. Each well of MyoTACTIC contains a cell seeding chamber and two vertical, flexible micro-posts that support hMMT self-organization and whose movement is tracked in short videos to measure hMMT force in situ. We used a combination of 3D printing and microfabrication techniques to produce a reusable PU mold that is employed to cast a nearly unlimited number of PDMS devices containing all 96-wells together with all well features. Single step casting eliminates the perceived challenge or technical issues arising in the use of reported culture platforms or cumbersome inserts containing vertical posts (Agrawal et al., 2017; Maffioletti et al., 2018; Powell et al., 2002; Vandenburgh et al., 2008, 2009). The MyoTACTIC platform makes hMMT production fast, reproducible, and user-friendly.

As mentioned, a notable advantage of MyoTACTIC is its capacity to quantify hMMT active force non-invasively and in situ, in a 96-well format by using post deflection captured in videos coupled with our Python tracking script. The non-invasive nature of our readouts are of critical importance since they enable longitudinal studies of drug effects on hMMTs at different time points. This challenge is faced by currently available methods where assessment of 3D tissue passive force (Agrawal et al., 2017), stimulated 2D culture contractility (Young et al., 2018), or stimulated 3D tissue force (Hinds et al., 2011; Madden et al., 2015) are implemented as endpoint assays. While there are certainly examples of 3D culture platforms that enable force measurement based on flexible-post deflection and image analysis (Osaki et al., 2018; Sakar et al., 2012; Vandenburgh et al., 2008, 2009), we expect that the 96-well footprint, one step fabrication of a stand-alone device, and ease of use is key to see widespread adoption by other researchers.

While 3D printing is an excellent technique for product prototyping due to its speed and cost-effectiveness, the technology still faces limitations with regards to the resolution of the technique for microfabrication. As seen in our supplementary data, we observed the presence of uneven surfaces behind each post in our first printing trials which prevented hMMTs from moving upward to the intended position just below the slanted region (**Supplementary Figure 2b**). We hypothesized that this was due to the 3D printing resin which remained intact despite extensive wax removal efforts. Consistently, modifying the printing direction improved the uneven surfaces, however, a certain level of uneven surface remained on micro-post structures which indicates the limitation of the technology for microscale printing. This fabrication challenge also serves to preclude the possibility of reporting hMMT absolute force measurements since hMMTs do not necessarily rise to the micro-post position where Microsquisher calibration is conducted (**Figure 3a-b**). However, given the reproducibility of micro-post mechanical measurements obtained across wells in a MyoTACTIC device (**Figure 3c**), we are confident in the overall design and its implementation to quantify changes in relative force.

Another point to consider is damage to the posts that occurs during fabrication Steps 1 to 3 (**Figure 1a**), which leads to the formation of unusable wells in the hard PU negative mold. In the course of our study, we noticed that the majority of wells containing a broken post were closest to the portion of the plate that was removed last in Step 3 (**Figure 1a**). We hypothesize that this is due to the sharp angles required for the final portion of plate demolding. In our hands, our best PU mold has 56 defect-free wells out of 96 total. Notably, after fabrication of the PU negative mold, Step 4 routinely yields many rounds of PDMS casting with no damage to the remaining wells. This validates the fitness of PU as the material of choice for PDMS microfabrication. To achieve defect-free MyoTACTIC devices, an alternative method to fabricate the final product after the prototyping stage, such as micromachining, could be employed to address the 3D printing shortcomings noted above.

We next conducted studies to confirm that the hMMTs cultured in MyoTACTIC exhibit similar structural and functional characteristics reported in previously published work (Bakooshli et al., 2018; Madden et al., 2015). In this regard, we show that MyoTACTIC enables the formation of hMMTs that display comparable muscle fiber hypertrophy, adult contractile protein expression, and calcium handling trends as reported in larger engineered human skeletal muscle tissue formats (**Figures 2** and **4**). Consistently, hMMTs cultured in MyoTACTIC exhibited formation of cross-striated muscle fibers, in as early as 7 days of differentiation, that contained clustered AChRs and responded to biochemical (ACh) and electrical stimuli by contracting (**Figures 2–4** and **Supplementary Figure 3c**). Using GCaMP6 transduced human skeletal muscle progenitor cells and capitalizing on the transparency of the MyoTACTIC, we recorded hMMT calcium transients in situ and over time from Day 7 to 14 and observed the maturation process (**Figure 4**).

To validate the application of MyoTACTIC to drug testing, we first focused our studies on compounds with well-studied effects on human skeletal muscle (Staffa et al., 2002; Tamraz et al., 2013; Velloso, 2008; Viguerie et al., 2012). An advantage of MyoTACTIC is the ability to support long-term culture. Hence, we focused all of our biological studies on compound administration regimes that started on Day 7 of culture when muscle fibers were competent to produce force (**Figure 3**) and few mononucleated cells remained (**Supplementary Figure 3b**), and assessed hMMTs one week later. As expected, dexamethasone and cerivastatin induced muscle fiber atrophy, as observed in histological analysis, and reduced hMMT contractile force generation, in a dose-dependent manner (**Figure 5**).

We next turned our attention to demonstrating the capacity of MyoTACTIC to detect compounds that increase strength. Studies in animals (Coleman et al., 1995; Musarò et al., 2001; Ye et al., 2013) support a role for IGF-1 in muscle hypertrophy. Culture studies in which IGF-1 is delivered to mono-nucleated myogenic progenitors when exposed to differentiation conditions in 2D culture (Jacquemin et al., 2004; Vandenburgh et al., 1991; Young et al., 2018), or to nascent myotubes in 2D (Rommel et al., 1999, 2001) or 3D (Vandenburgh et al., 2008b) culture also observed hypertrophic effects by immunohistological analysis. Interestingly, we did not find a change in the width of IGF-1 treated hMMT muscle fibers even at a dose reported by others to induce muscle fiber hypertrophy (Rommel et al., 2001). We expect this is explained by a difference in treatment regime, with other studies initiating IGF-1 treatment regimens at earlier stages of differentiation. Regardless, we observed a significant increase in the force generated by IGF-1 treated hMMTs, which might be attributed to IGF-1 effects on skeletal muscle fiber protein translation (Bodine et al., 2001; Rommel et al., 2001). Indeed, visual inspection of IGF-1 treated fibers reveals a uniformity in fiber width along the entire structure hinting at contractile apparatus maturation. This signifies the predictive value of the MyoTACTIC platform in capturing drug and biomolecule effects using a clinically relevant functional readout for skeletal muscle.

hMMT studies offer the unique opportunity to decouple direct and indirect effects of systemic treatments on skeletal muscle health. To highlight this point, we sought to interrogate potential direct influences of chemotherapeutic agents on hMMT morphology and strength (**Figure 6**). Cachexia, a specific type of muscle wasting, is frequently described in cancer patients, including those with pancreatic cancer (Argilés et al., 2014; Dodson et al., 2011; Sun et al., 2015), and it is associated with decreased life expectancy (Dimitriu et al., 2005). The cause of cancer-associated cachexia is a very active area of research.

In general, the direct effect of cancer chemotherapeutic agents on skeletal muscle health is understudied, since most are developed with a mitosis-targeting mechanism of action, and skeletal muscle fibers are post-mitotic. However, evidence suggested that human skeletal muscle might be a direct target of two mitosis-targeting chemotherapeutic drugs; Gemcitabine and Irinotecan. First, a study in healthy mice evaluated skeletal muscle following systemic treatment with Folfox or Folfiri, two commonly implemented pancreatic cancer patient chemotherapy cocktails. Loss of muscle mass and weakness was observed in animals treated with Folfiri and not Folfox, with Irinotecan being the variable discerning the two cocktails, though this point was not explored in the report (Barreto et al., 2016). Interestingly, a study of mouse C2C12 cells in culture noted that Irinotecan induced changes in cycling myoblast mitochondrial activity, but myotube treatment had little influence on width (Rybalka et al., 2018). We implemented MyoTACTIC to study the direct effects of these widely used chemotherapeutic agents on hMMT function. Strikingly, a one-time treatment of a clinical dose of Irinotecan (16 μM, (Chabot, 1997) on Day 7 led to a significant decrease in hMMT contractile force as compared to carrier control treatment. Most notably, histologically apparent effects on muscle fiber integrity were only uncovered at a higher dose of Irinotecan (72 μM), underlying the sensitivity of our non-invasive functional assay. Even the highest dose of Gemcitabine produced no detectable effect on muscle fiber morphology or strength. Together, this data provides motivation to conduct epidemiological studies in this area with the goal of informing chemotherapeutic regimes tailored to the patient as their needs evolve over the course of treatment.

Together, our data highlight the ease-of-use, reproducibility, robustness, and versatility of MyoTACTIC, a platform created for in vitro human skeletal muscle drug testing.

## METHODS

### Platform fabrication for human muscle micro-tissue (hMMT) culture

Fabrication of the final polydimethylsiloxane (PDMS) platform (MyoTACTIC) was accomplished through sequential fabrication of intermediate plates as shown in **Figure 1a**. The original MyoTACTIC template was designed in SOLIDWORKS (**Supplementary Figure 1**) and 3D-printed using ProJet MJP 3500 Series from 3D SYSTEMS. VisiJet M3 Crystal was used as the printing material. The template was then cast into a flexible PDMS plate using Sylgard 184 silicone elastomer kit (Dow Corning) (**Figure 1a**, step 1). During casting, the template was lightly sprayed with Ease Release^®^ 200 release agent (Smooth-On) and left in a chemical hood for 10 to 15 minutes to ensure consistent spread of the release agent. Liquid PDMS (10 parts monomer to one part curing agent) was degassed in a vacuum chamber and poured onto the original 3D-printed template. A container was used to keep the liquid PDMS in place. Next, the container (including the template and the liquid PDMS) was degassed in a vacuum chamber for 45 minutes to remove any bubbles trapped within the PDMS. The container was then placed in an isothermal oven set to 60 °C overnight. The next day the cured negative PDMS mold was manually separated from the template. The negative PDMS plate was then used as a template to cast a positive PDMS plate (**Figure 1a**, step 2). To ensure the release of the second PDMS part from the negative PDMS mold during this PDMS to PDMS casting step, a surface modification was applied to the template as previously described (Shao et al., 2012). Briefly, the negative PDMS mold was plasma-treated for 2 minutes in a plasma chamber (Harrick Plasma). Plasma-treated negative PDMS mold was immediately coated with 1H, 1H, 2H, 2H-perfluorodecyltrichlorosilane (Sigma) in a vacuum chamber for at least 3 hours. Next, liquid PDMS (10 parts monomer to one part curing agent) was poured onto the silane-coated PDMS negative. To promote the penetration of liquid PDMS into the small features, the negative mold was placed inside a vacuum chamber and thorough degassing was performed for at least one hour with intermittent vacuum breaks. Next the container was placed in an isothermal oven at 60 °C overnight. The cured positive PDMS mold was then separated from the negative PDMS mold. Next, the positive PDMS mold was used as a template to fabricate a negative polyurethane (PU) plate (**Figure 1a**, step 3). To cast the negative PU plate, the positive PDMS plate was sprayed with a light layer of Ease Release^®^ 200 and left at room temperature in a chemical safety hood for at least 15 minutes to ensure even spread of the release agent. Next, Smooth-Cast^®^ 310 liquid plastic (Smooth-On) was prepared based on the manufacturer’s instructions and was poured on top of the positive PDMS mold in a container and left at room temperature until fully solidified (minimum of 3 hours). Subsequently the PDMS plate was separated from the PU mold. At this stage, the rigid PU negative mold enabled one-step fabrication of the MyoTACTIC platform. Copies of MyoTACTIC were then produced by repeatedly using the PU negative mold as a template (**Figure 1a**, step 4). The PU template was sprayed with a light layer of Ease Release^®^ 200 and liquid PDMS (15 parts monomer to one part curing agent) was poured onto the PU mold. Next, the mold was degassed thoroughly in a vacuum chamber to ensure the total penetration of PDMS into the small features of the PU mold. The construct was placed in an isothermal oven at 60 °C for at least 3 hours, after which the final PDMS device, MyoTACTIC, (**Figure 1b**) was separated from the negative PU mold.

### Quantification of micro-post mechanical properties

Mechanical properties of the micro-posts were determined using a Microsquisher device (CellScale). Briefly, micro-posts with their PDMS base were excised from each well of a MyoTACTIC using a sharp blade and mounted in the Microsquisher test chamber, immersed in 37 °C PBS. A micro-wire (**Figure 3a**) connected to a force transducer was used to deflect microposts with known amounts of force (**Supplementary Movie S15**), and quantified the extent of micro-post displacement that was induced. In this way, Microsquisher analysis quantified the relationship between the magnitude of force applied to micro-posts, and their displacement (**Figure 3b-c**).

### Human skeletal muscle biopsy material

Human skeletal muscle tissues removed during scheduled surgical procedures and designated for disposal were utilized in this study in accordance with St. Michael’s Hospital research ethics board and University of Toronto administrative ethics review approval. Small skeletal muscle samples (~1 cm^3^) were obtained from the multifidus muscle of patients undergoing lumbar spine surgery.

### Primary human myoblast preparation and expansion

Primary human myoblast cell lines were established and maintained as previously described (Bakooshli et al., 2018). Briefly, human skeletal muscle samples were minced and then dissociated into a single cell slurry with Clostridium histolyticum collagenase (Sigma, 630 U/mL) and dispase (Roche, 0.03 U/mL) in Dulbecco’s Modified Eagle’s medium (DMEM; Gibco). The cell suspension was passed multiple times through a 20G needle to facilitate the release of the mononucleated cell population and subsequently depleted of red blood cells with a brief incubation in red blood cell lysis buffer (**Table S2**). The resulting cell suspension containing a mixed population of myoblasts and fibroblast-like cells was plated in a collagen-coated tissue culture dish containing myoblast growth medium: F-10 media (Life Technologies), 20% fetal bovine serum (Gibco), 5 ng/mL basic fibroblast growth factor (bFGF; ImmunoTools) and 1% penicillin-streptomycin (Life Technologies). After one passage, the cell culture mixture was stained with an antibody recognizing the neural cell adhesion molecule (NCAM/CD56; BD Pharmingen; **Table S1**), and the myogenic progenitor (CD56+) population was separated and purified using fluorescence-activated cell sorting (FACS) and maintained on collagen-coated dishes in the myoblast growth medium. Subsequent experiments utilized low passage cultures (P4 - P9).

### Fabrication of human muscle microtissues (hMMTs)

Human muscle microtissues were generated following our previously described method (Bakooshli et al., 2018). Briefly, FACS purified CD56^+^ myogenic progenitor cells were suspended in a hydrogel mixture (**Table S2**) at 15 × 10^6^ cells/ml. Thrombin (Sigma) was added at 0.2 to 0.5 unit per mg of fibrinogen prior to seeding the cell/hydrogel mixture in MyoTACTIC wells. Tissues were then incubated for 5 minutes at 37 °C to expedite thrombin-mediated fibrin polymerization. Myoblast growth media (**Table S2**) lacking bFGF, but containing 1.5 mg/mL 6-aminocaproic acid (ACA; Sigma), was added. 2 days later the growth media was exchanged to myoblast differentiation medium (**Table S2**) containing 2 mg/ml ACA. This time point is referred to as differentiation Day 0. Half of the culture media was exchanged every other day thereafter until the desired experimental endpoint.

### hMMTs drug treatments

Drugs were purchased as listed in **Table S3**. All drugs were sterile-filtered for use before adding to the media. Dexamethasone and Cerivastatin were dissolved in ethanol and ddH2O at 2.548 mM and 4.352 mM respectively. Subsequently, 50X stock solutions were prepared for each dose by diluting the drug solutions in DMEM. IGF-1 was prepared in 10 mM HCl solution at 100 μM of the highest dose and diluted to 50X stock solutions for each dose using DMEM. Gemcitabine was reconstituted in PBS at 40X (12.8 mM) stock solution for its highest dose and diluted in DMEM for lower doses. Irinotecan was prepared in DMSO at 31.75 mM of its highest dose and diluted to 21X stock solutions for each dose using DMEM. Vehicle solutions (ethanol, ddH2O, PBS, HCl, or DMSO) were added to the culture media of control hMMTs at concentrations corresponding to the highest drug doses administered. Drug treatment of hMMTs was initiated on differentiation Day 7. To maintain the concentration of drugs in the media, Dexamethasone and Cerivastatin were added to the hMMT media every other day during media change time points, while IGF-1 was added daily at full dose. Chemotherapeutic drugs were added to the hMMT media once on differentiation Day 7. Half the media was replaced with fresh media every other day thereafter until differentiation Day 14.

### Immunostaining and fluorescence microscopy

hMMTs were fixed and labeled for confocal imaging as previously described (Bakooshli et al., 2018). Briefly, hMMTs were fixed on the posts in 4% PFA for 15 minutes and then washed with PBS. Following fixation, samples were incubated in blocking solution (**Table S2**) for 1 hour at room temperature or overnight at 4 °C. Samples were then incubated in primary antibody (**Table S1**) solutions diluted in blocking solution (**Table S2**) overnight at 4 °C. After several washes in blocking solution or PBS, samples were incubated with appropriate secondary antibodies diluted in the blocking solution for 30-60 minutes at room temperature. Hoechst 33342 or DRAQ5 (ThermoFisher) were used to counterstain cell nuclei. Confocal images were acquired with Fluoview-10 software using an Olympus IX83 inverted microscope. Phase-contrast images were acquired with CellSense™ software using an Olympus IX83 microscope equipped with an Olympus DP80 dual CCD color and monochrome camera or an Apple^®^ iPhone^®^ SE. Images were analyzed and prepared for publication using NIH ImageJ software.

### Myofiber size analysis

Myofiber size was analyzed as previously described (Bakooshli et al., 2018). Briefly, myofiber size was measured by assessing 20X and 40X magnification confocal images of immunostained hMMTs for sarcomeric α-actinin. Z-stack images of hMMTs were analyzed to quantify the diameter of each intact muscle fiber at the center of each image using NIH ImageJ. Average myofiber diameter was determined per image, and these values were then averaged to calculate the average myofiber diameter for each hMMT.

### Western blotting

hMMTs were collected at the mentioned time points and immediately lysed in RIPA buffer (Thermofisher) containing Halt™ protease inhibitor cocktail (Thermofisher). Total protein concentration was measured using BCA assay kit (Thermofisher) and 20 to 25 μg of protein was run on an 8% SDS-PAGE gel. Western blot was performed as described previously (Bakooshli et al., 2018).

### hMMT electrical stimulation

Two electrodes (25G x 5/8 BD™ Regular Bevel Needles) were inserted behind the posts into each MyoTACTIC well to generate an electric field parallel to the fibers. Electrodes were sterilized using 70% ethanol before insertion into wells. Electrodes were then connected to a commercial pulse generator (Rigol DG1022U), using nickel coated copper wires (McMaster-Carr) and alligator clamps. A Rigol DS1102E digital oscilloscope was used to confirm the frequency and amplitude of the signals before connecting the pulse generator to the wires. hMMTs were stimulated using square pulses with 20% duty cycle, 5V amplitude (electrical field strength of 10 V/cm), and 0.5 Hz and 20 Hz frequency for twitch and tetanus contractions, respectively. A graphical description of the electrical stimulation setup is presented in **Supplementary Figure 4**.

### hMMT biochemical stimulation

Acetylcholine was reconstituted to produce a 200 mM stock solution in DMEM (100X) and was diluted to the final working concentration by addition directly into each MyoTACTIC wells containing hMMTs.

### hMMT calcium transient analysis

Human myogenic progenitor cells expressing the MHCK7-GCaMP6 reporter were generated as previously described (Bakooshli et al., 2018; Madden et al., 2015). hMMTs expressing GCaMP6 were imaged using an Olympus IX83 microscope at different time points following differentiation. Movies were recorded at 4X magnification at 11 frames per second using an Olympus DP80 dual CCD color and monochrome camera with a FITC filter and CellSense™ software. MyoTACTIC wells containing hMMTs were placed on the microscope stage equipped with temperature and gas modules to simulate physiological conditions (37 °C and 5% CO2) during data collection. A region of interest in the center of each hMMT was defined for fluorescence analysis (**Supplementary Movie S16**) and movies were then analyzed using NIH ImageJ software. Relative changes in fluorescence signal were measured and are reported as ΔF/F_0_ = (F_immediate_ − F_baseline_)/(F_baseline_). The relative change in fluorescence signal was plotted against time and the peak signal of 5 to 8 consecutive stimulations (contractions) were averaged for each hMMT.

### Measurement of hMMT contractile force

Movies of post deflection were captured using an Apple^®^ iPhone^®^ SE during electrical stimulation of hMMTs under 10X magnification using an Olympus IX83 microscope and a LabCam™ iPhone microscope mount (**Supplementary Figure 4c**). Movies were then analyzed using a custom-written Python computer vision script (described in the **Appendix**), which determined post displacement in pixels during post deflections. hMMTs were induced to contract 5 times, and the maximum post displacements were averaged to calculate the average post deflection per hMMT.

### EdU analysis

Click-iT^®^ EdU cell proliferation assay kit (Thermofisher) was used for cell cycle analysis. 5-ethynyl-2’-deoxyuridine (EdU) was added at 10 μM to the culture medium of hMMTs on Day 0, Day 1, or Day 7 of differentiation (**Supplementary Figure 3b**). 19 hours post-EdU administration, hMMTs were fixed and stained following the user manual. Briefly, fixed hMMTs were permeabilized in 0.3% Triton X-100 solution for a minimum of 15 minutes. Next, cells were counterstained for Alexa Fluor 647 using anti-EdU reaction solution for 30 minutes at room temperature. Nuclei were counterstained with Hoechst 33342. Images were acquired using confocal microscopy as mentioned before.

### Statistical analysis

hMMT-level data such as average myofiber diameter per hMMT, average ΔF/F_0_ per hMMT, and average post deflection per hMMT, constituted technical replicates. Technical replicate data from hMMTs seeded at the same time and from the same cell source (i.e. from the same biological replicate) were averaged together to calculate a single biological replicate average. All hMMT experiments were performed with a minimum of 3 biological replicates. Statistical testing for significance was conducted on these biological replicate averages, while technical replicate data were used to verify the parametric assumptions of residual normality and homogeneity of variance via the Shapiro-Wilk test and the Brown-Forsythe test, respectively. Technical replicate data were transformed to pass the parametric assumptions when necessary, whereupon the relevant statistical test was performed on similarly transformed biological replicate data. When transformation of data did not resolve violations of the parametric assumptions, non-parametric tests were employed. Parametric tests used included the one-way ANOVA followed by Tukey’s multiple comparison test, the independent student’s t-test, and the independent student’s t-test with Welch’s correction. Non-parametric tests used included the Kruskal-Wallis test followed by Dunn’s multiple comparisons test. Statistical analyses were completed using GraphPad Prism 6 (La Jolla, USA).

## Acknowledgements

We would also like to acknowledge the following sources for funding this study: NSERC CREATE TOeP to M.E.A., H.Y.A., M.A.B. S.D.; Ontario Graduate Scholarship to H.Y.A., M.A.B; Toronto Musculoskeletal Centre and Krembil Foundation to M.A.B.; and Ontario Research Fund (31390), Canada Research Chair Program (950-231201), Canada Foundation for Innovation (31390), Canada First Research Excellence Fund ‘Medicine by Design’, the Ontario Institute for Regenerative Medicine, Natural Sciences and Engineering Research Council (RGPIN 435724-13) to P.M.G.

## Author contributions

MEA, HYA, and MAB designed and performed the experiments, analyzed and interpreted the data, and prepared the manuscript. SD and NT were involved in and facilitated culture device design and fabrication, and prepared the manuscript. KT consented study tissue donors and facilitated transfer of samples. HA and HG surgically resected and provided skeletal muscle biopsies, interpreted the data, and revised the manuscript. PZ and PMG designed the experiments, supervised the work, interpreted the data, prepared and revised the manuscript.

## Materials & Correspondence

Correspondence should be addressed to Penney M Gilbert.

## Supplemental Information

### Supplemental Figures and Figure Legends

#### Link to Supplemental Figures

https://www.dropbox.com/sh/uvrhe85y5wu1bez/AAAqc-wutyH4ji5DVxrCtUTZa?dl=0

### Supplemental Figure Legends

**Supplemental Figure 1. MyoTACTIC three-dimensional computer aided design drawing**.

**(a)** Three-dimensional computer aided design (3D CAD) drawing of MyoTACTIC master plate. Dimensions are in mm. **(b)** Detailed drawings of the individual wells of MyoTACTIC into which the hMMTs are seeded in. Dimensions are in mm. **(c)** Total volume and surface area of the upper and lower wells, and the individual posts in each MyoTACTIC well.

**Supplemental Figure 2. Primary human muscle progenitor enrichment and hMMT tissue formation in MyoTACTIC. (a)** Representative fluorescence activated cell sorting (FACS) dot plot of primary human myoblasts (CD56+) sorted after one passage from the mix culture preparation from a patient muscle. **(b)** Surface roughness on the outer side of the micro-posts from the 3D printed platform (left panel) results in inconsistent vertical positioning of the hMMTs post remodeling as shown on the right panel. Scale bar 1 mm. Posts are outlined with yellow dashed lines on the right panel.

**Supplemental Figure 3. Characterization of hMMTs produced in MyoTACTIC. (a)** Quantification of hMMT width over the course of culture time. n= minimum of 16 hMMTs from 3 muscle patient donors per time point. Data are presented as mean ± SEM. **(b)** Representative stitched confocal images of hMMTs counter stained with Hoechst to label nuclei (blue) and labeled for EdU (green) to visualize proliferating cells at days 0 (top panel), 1 (middle panel), and 7 (bottom panel) differentiation. hMMTs are outlined in white dashed lines and location of micro-posts are outlined with yellow dashed circles. Scale bar 500 μm. **(c)** Representative confocal images of the hMMTs on day 7, 10, and 14 of differentiation immunostained for sarcomeric a-actinin (SAA, red) and a-bungarotoxin (BTX; green) to label AChR clusters. Scale bar 50 μm. **(d)** Left panel: Schematic of 3 different media change regimens and their effect on hMMT myofiber diameter. + represents replacement of all of the culture media with fresh differentiation media, * indicates replacement of half of the culture media with fresh differentiation media. Right panel: Dot plot quantification of the average myofiber diameter of hMMTs treated with the regimens presented in the left panel, on day 14 of differentiation. **p < 0.01. Values are reported as mean ± SEM. Significance was determined by one-way ANOVA followed by multiple comparisons to compare differences between groups using Tukey’s multiple comparisons test.

**Supplemental Figure 4. Configuration of the electrical stimulation setup. (a)** Olympus IX83 inverted microscope equipped with a fluorescence lamp was used to capture movies during electrical stimulation for post displacements and calcium handling of hMMTs. Platinum wires were hooked up to a commercial function generator (Rigol DG 1022U, outlined in yellow dashed circle on bottom right). An Apple^®^ iPhone^®^ SE camera in combination with a LabCam™ mount was used to capture the movies (outlined in yellow dashed box in the middle). hMMTs were kept under physiological conditions using a stage top mini incubator equipped with temperature and gas modules to control the CO2 concentrations. **(b)** A Rigol DS1102E digital oscilloscope was used to confirm the frequency and amplitude of signals before connecting the pulse generator to the platinum wires. **(c)** Representative photo of the Apple^®^ iPhone^®^ SE and the LabCam™ mounted on the eyepiece of the Olympus IX83 microscope. **(d)** Representative picture indicating the location of the electrodes during the electrical stimulations (Xs on the left and right sides of the hMMT).

### Supplemental Movies and Movie Captions

#### Link to Supplemental Movies

https://www.dropbox.com/sh/uvrhe85y5wu1bez/AAAqc-wutyH4ji5DVxrCtUTZa?dl=0

### Supplemental Movie Captions

**Movie S1. hMMT spontaneous contractions on Day 10 of differentiation**. A series of three representative bright-field videos of hMMTs after 10 days of differentiation exhibiting spontaneous contractions. Movies were recorded using an Apple^®^ iPhone^®^ SE.

**Movie S2. hMMTs generate twitch contractions in response to low frequency (0.5 Hz) electrical stimuli**. A representative bright-field video of a hMMT generating a twitch contraction in response to low frequency (0.5 Hz) electrical stimuli. Movie was recorded using an Apple^®^ iPhone^®^ SE under 4X magnification.

**Movie S3. hMMTs generate tetanus contractions in response to high frequency (20 Hz) electrical stimuli**. A representative bright-field video of a hMMT generating a tetanus contraction in response to high frequency (20 Hz) electrical stimuli. Movie was recorded using an Apple^®^ iPhone^®^ SE under 4X.

**Movie S4. Measurement of micro-post deflection in response to 20 Hz electrical stimulation of hMMTs using a custom Python computer vision script**. A representative bright-field video of a micro-post deflection being tracked by the custom-written Python computer vision script (blue box) during 5 tetanus contractions. hMMT is stimulated using high frequency electrical stimuli (20Hz) to generate tetanus contractions. Movie was recorded using an Apple^®^ iPhone^®^ SE under 10X magnification and is 10X fast forwarded.

**Movie S5. Micro-post deflection in response to 20 Hz electrical stimulation of hMMTs on Day 7, 10, and 14 of differentiation**. A series of three bright-field representative movies of micro-post deflection in response to hMMT tetanus contractions on Days 7, 10, and 14 of differentiation. hMMTs are stimulated using high frequency electrical stimuli (20Hz) to generate tetanus contractions. Movies were recorded using an Apple^®^ iPhone^®^ SE under 4X magnification and are 3X fast forwarded.

**Movie S6. Spontaneous calcium handling of hMMTs on Day 7 differentiation**. A representative epifluorescence time-lapse video of hMMTs after 7 days of culture demonstrating spontaneous calcium transients. Calcium transients are visualized in green by following the GCaMP6 calcium reporter signal that was transduced into the human muscle cells.

**Movie S7. hMMT twitch and tetanus contraction calcium handling on differentiation Days 7, 10, and 14**. A series of three representative videos of hMMTs after 7, 10, and 14 days of differentiation stimulated with 0.5 Hz electrical stimuli. Muscle fiber calcium transients are visualized in green by following the GCaMP6 calcium reporter signal that was transduced into the human muscle cells. Movies are fast forwarded 3X.

**Movie S8. hMMT calcium handling on differentiation Days 7, 10, and 14 in response to biochemical stimuli**. A series of three representative videos of hMMTs after 7, 10, and 14 days of differentiation and stimulated with ACh (2mM). Muscle fiber calcium transients are visualized in green by following a GCaMP6 calcium reporter signal that was transduced into the human muscle cells. Movies are fast forwarded 3X.

**Movie S9. d-tubocurarine treatment blocks hMMT response to ACh but has no effect on hMMT response to electrical stimuli**.

A series of three representative videos of a hMMTs on Day 14 of differentiation following treatment with d-tubocurarine (25 μM) and stimulation with electrical (0.5 Hz and 20Hz) and biochemical (ACh, 2mM) stimuli. Muscle fiber calcium transients are visualized in green by following a GCaMP6 calcium reporter that was transduced into the human muscle cells. Movies are fast forwarded 3X.

**Movie S10. Dexamethasone treatment reduces the contractile force generation of hMMTs compared to control (untreated) hMMTs as indicated by post deflection analysis**. A series of two bright-field representative movies of micro-post deflection in response to hMMT tetanus contractions on Day 14 of differentiation. hMMTs are treated with vehicle (DMSO, control) or Dexamethasone (10 nM) from Day 7 to day 14 of differentiation. hMMTs are stimulated using high frequency electrical stimuli (20Hz) to generate tetanus contractions. Movies were recorded using an Apple^®^ iPhone^®^ SE under 4X magnification and are 6X fast forwarded.

**Movie S11. Cerivastatin treatment reduces the contractile force generation of hMMTs compared to control (untreated) hMMTs as indicated by post deflection analysis**. A series of two bright-field representative movies of micro-post deflection in response to hMMT tetanus contractions on Day 14 of differentiation. hMMTs are treated with vehicle (DMSO, control) or Cerivastatin (10 nM) from Day 7 to 14 of differentiation. hMMTs are stimulated using high frequency electrical stimuli (20Hz) to generate tetanus contractions. Movies were recorded using an Apple^®^ iPhone^®^ SE under 10X magnification and are 6X fast forwarded.

**Movie S12. IGF-1 treatment increases the contractile force generation of hMMTs compared to control (untreated) hMMTs as indicated by post deflection analysis**. A series of two bright-field representative movies of micro-post deflection in response to hMMT tetanus contractions on Day 14 of differentiation. hMMTs are treated with vehicle (DMEM, control) or IGF-1 (100 nM) from Day 7 to 14 of differentiation. hMMTs are stimulated using high frequency electrical stimuli (20Hz) to generate tetanus contractions. Movies were recorded using an Apple^®^ iPhone^®^ SE under 10X magnification and are 6X fast forwarded.

**Movie S13. Gemcitabine treatment of hMMTs at supraphysiological dose does not affect their contractile force generation compared to control (untreated) hMMTs**. A series of two bright-field representative movies of micro-post deflection in response to hMMT tetanus contractions on Day 14 of differentiation. hMMTs are treated with one-time dose of vehicle (DMSO, control) or Gemcitabine (320 nM) on Day 7 of differentiation. hMMTs are stimulated using high frequency electrical stimuli (20Hz) to generate tetanus contractions. Movies were recorded using an Apple^®^ iPhone^®^ SE under 10X magnification and are 6X fast forwarded.

**Movie S14. Irinotecan treatment reduces the contractile force generation of hMMTs compared to control (untreated) hMMTs as indicated by post deflection analysis**. A series of three bright-field representative movies of micro-post deflection in response to hMMT tetanus contractions on Day 14 of differentiation. hMMTs are treated with one-time dose of vehicle (DMSO, control) or Irinotecan (16 nM, 72 nM) on Day 7 of differentiation. hMMTs are stimulated using high frequency electrical stimuli (20Hz) to generate tetanus contractions. Movies were recorded using an Apple^®^ iPhone^®^ SE under 10X magnification and are 6X fast forwarded.

**Movie S15. Measurement of force-displacement relation of MyoTACTIC micro-posts using Microsquisher**. Representative movie demonstrating the displacement of the microposts in response to the force exerted by the micro-wire connected to a force transduce to generate a force-displacement correlation for micro-post deflection. Location of the post and the micro-wire are indicated with black arrows.

**Movie S16. ROIs for calcium transients and contractile force analyses**. A bright-field movie of a hMMT under high frequency electrical stimulation (20Hz) captured with an Apple^®^ iPhone^®^ SE demonstrating various regions of interests for data collection and analysis.

**Table S1.** List of primary antibodies

| # | Antibody | Species | Dilution | Source |
| --- | --- | --- | --- | --- |
| 1 | Alexa Fluor 647 mouse<br>anti-human CD56 | Mouse | 1:20 | BD<br>Pharmingen |
| 2 | Anti- $\beta$ -tubulin | Rabbit | 1:5000 | Cell Signaling |
| 3 | DRAQ5 | - | 1:1000 | ThermoFisher |
| 4 | Hoechst 33342 | - | 1:1000 | ThermoFisher |
| 6 | Myosin heavy chain - fast | Mouse | 1:50 | DSHB |
| 7 | Myosin heavy chain - slow | Mouse | 1:50 | DSHB |
| 8 | Sarcomeric α-actinin | Mouse | 1:200 | Sigma |
| 9 | α-Bungarotoxin, Alexa<br>Fluor 647 conjugate | - | 1:500 | ThermoFisher |

**Table S2.** Cell Culture Media and Solutions

| # | Name | Details |
| --- | --- | --- |
| 1 | Blocking solution | 20 % goat serum, 0.3 % Triton-X 100 in PBS |
| 2 | Fibrinogen stock solution | 10 mg / mL fibrinogen in 0.9 % (wt / v)<br>NaCl solution in water |
| 3 | Human myoblast differentiation media | Dulbecco's Modified Eagle's medium (DMEM), 2 %<br>horse serum, 10 µg / mL insulin,<br>1% penicillin-streptomycin |
| 4 | Human myoblast growth media | Ham's F-10 nutrient mix, 20 % fetal bovine serum, 5 ng /<br>mL basic fibroblast growth factor,<br>1 % penicillin-streptomycin |
| 5 | Hydrogel mixture | Dulbecco's Modified Eagle's medium (DMEM) (40% v/v),<br>4 mg / mL bovine fibrinogen (40% v/v), Geltrex™ (20%<br>v/v), thrombin (0.2 unit / mg fibrinogen) |
| 6 | Milk based blocking solution | 5 % (wt / v) skim milk (BioShop) in TBST |
| 7 | Red blood cell lysis buffer | 15.5 mM NH<sub>4</sub>Cl, 1 mM KHCO<sub>3</sub>, 10 µM EDTA |
| 8 | Tris-buffered saline Tween (TBST) | 50 mM Tris (BioShop), 150 mM NaCl (Sigma),<br>0.1 % (v / v) Tween 20 (BioShop) |

**Table S3.** List of drugs

| **#** | **Name** | **Vendor (Product #)** |
| --- | --- | --- |
| 1 | Cerivastatin | Sigma (SML0005) |
| 2 | Dexamethasone | Sigma (D1756) |
| 3 | Gemcitabine | Sigma (G6423) |
| 4 | IGF-1 | Sigma (I1271) |
| 5 | Irinotecan | Cayman Chemical Company (14180) |

## Appendix Post tracking Python code

This computer vision script, when run, prompts the user to select a file containing the video of interest and select the region of interest on the MyoTACTIC post. Then, the script tracks the horizontal position of the post in pixels over the course of the video and calculates the displacements associated with human muscle microtissue contractions. For the experiments described here, the MyoTACTIC post edge farthest from the tissue was chosen as the region of interest. However other users may find different regions of interest to be more optimal for their own purposes. This post tracking script is written in Python 3, and requires the matplotlib library and the opencv-contrib library to run. This script has been tested on Windows 10 Home and macOS Version 10.14.2 operating systems.of the frames that the tracking is done on.

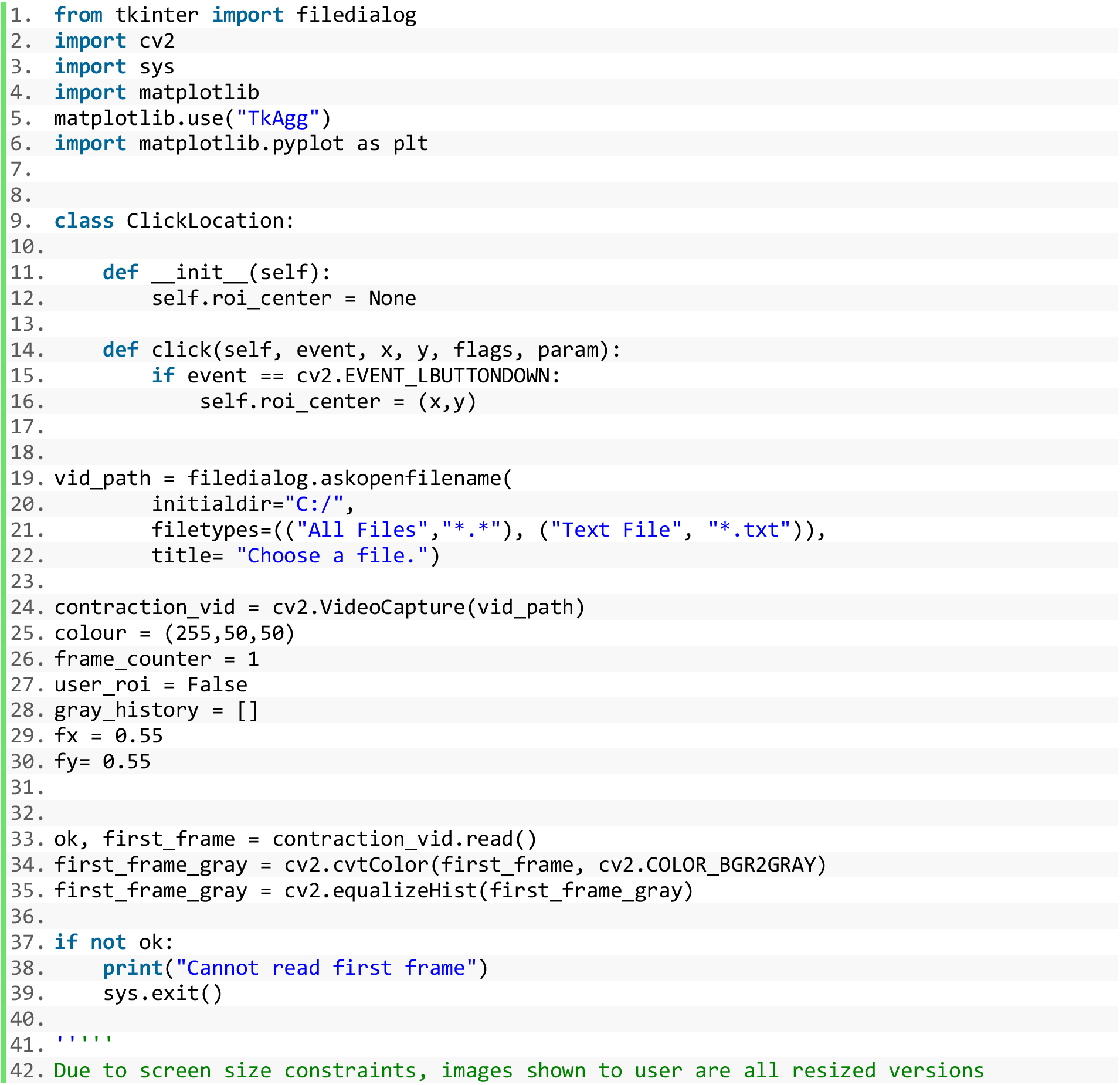
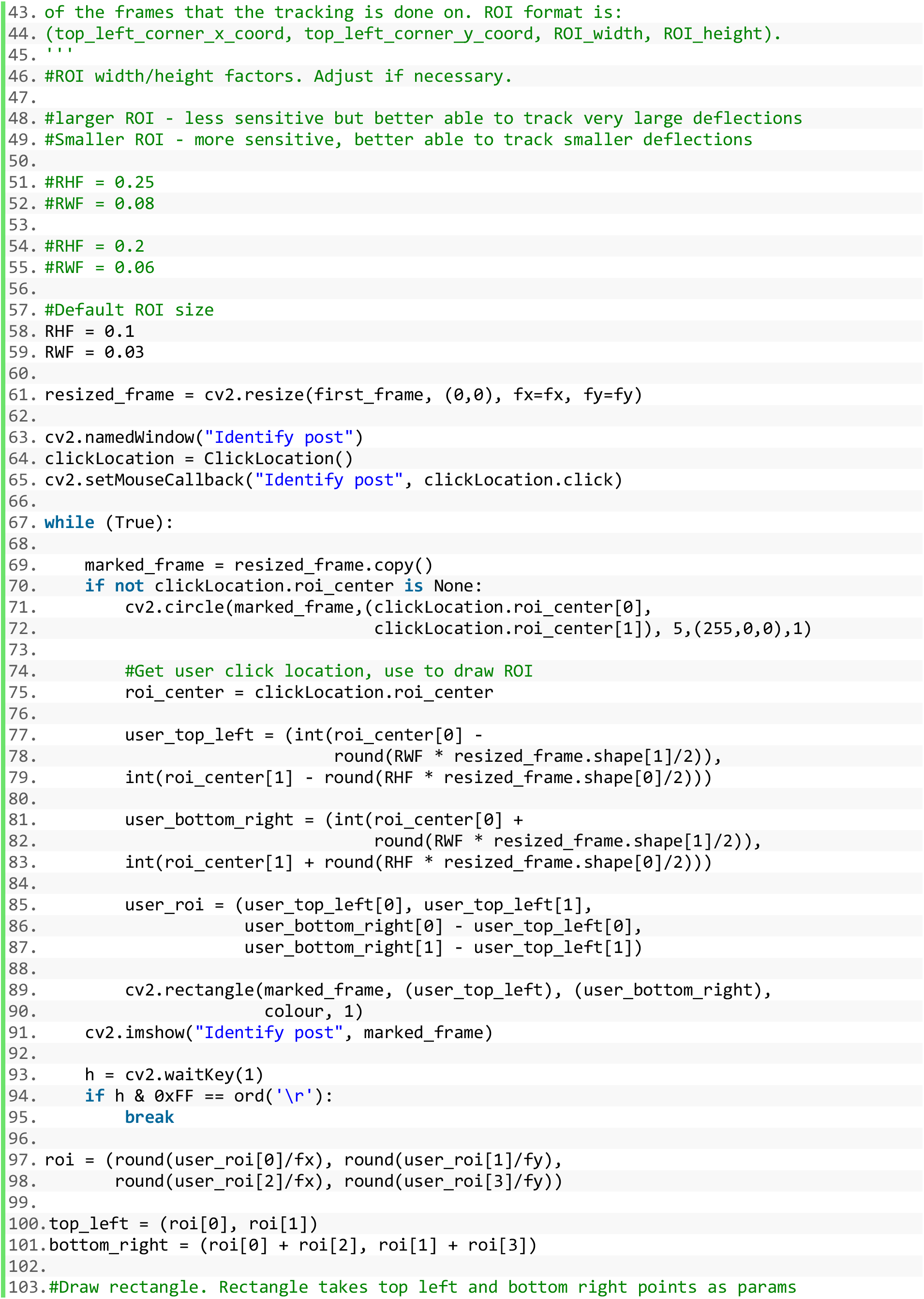
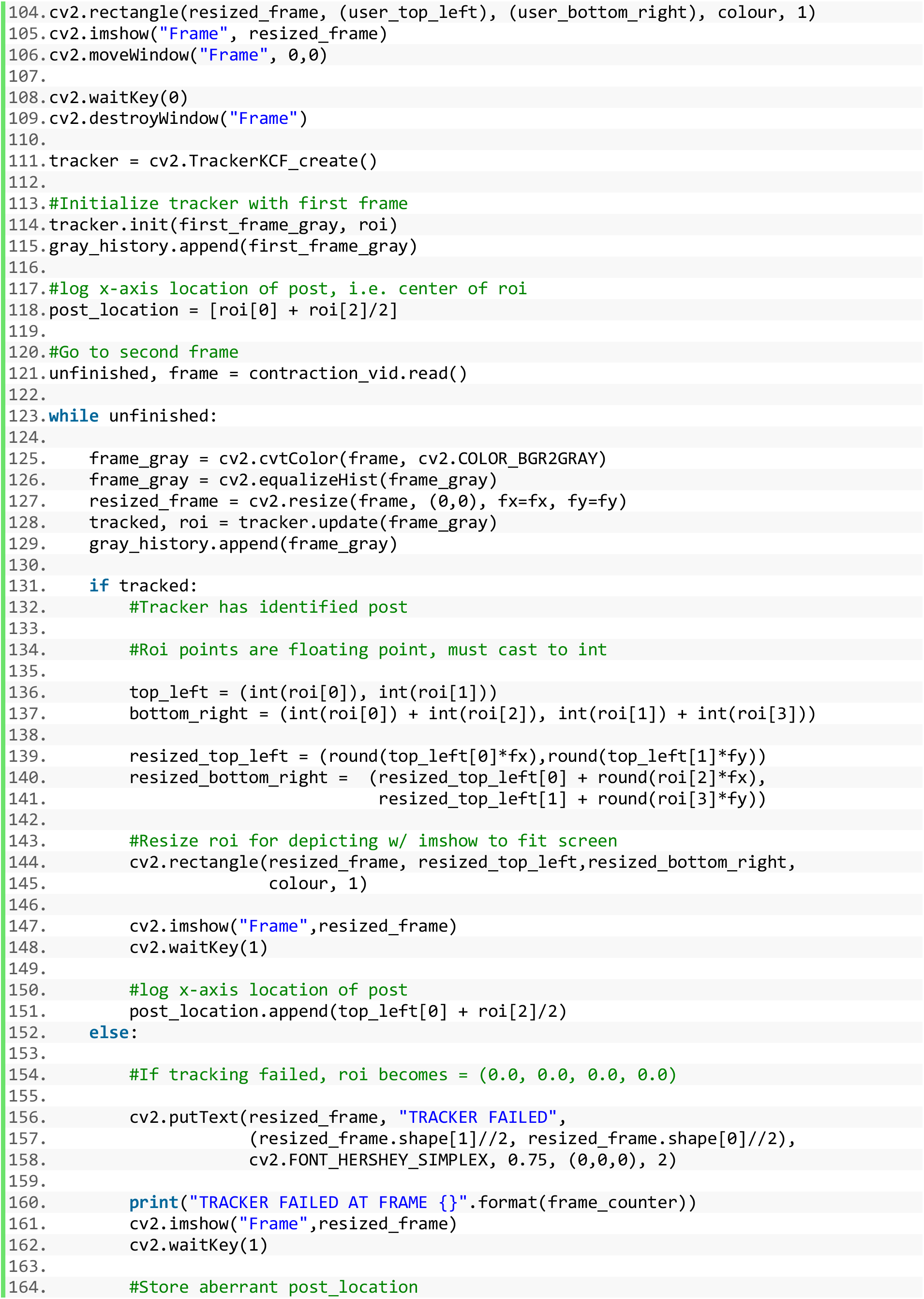
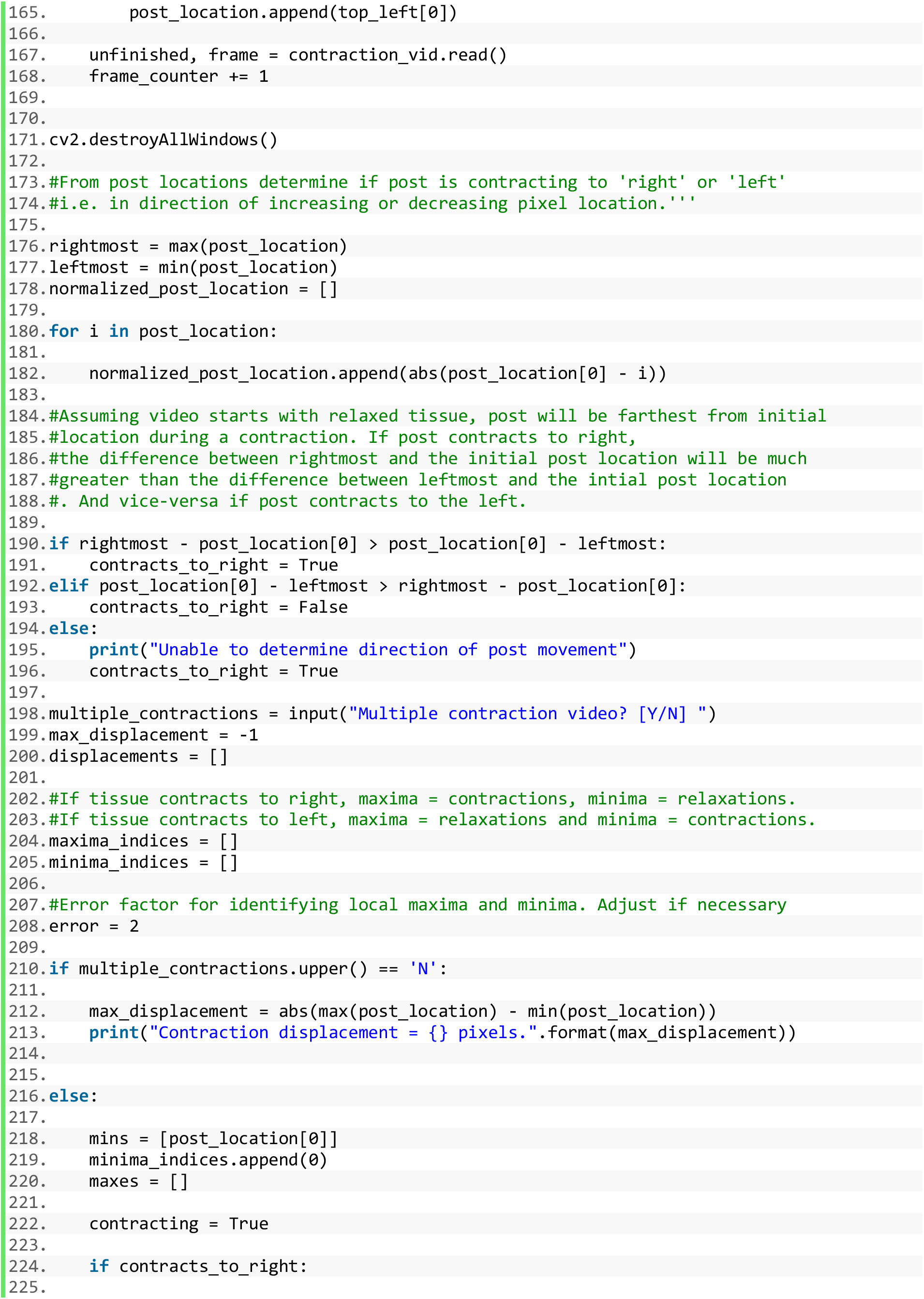
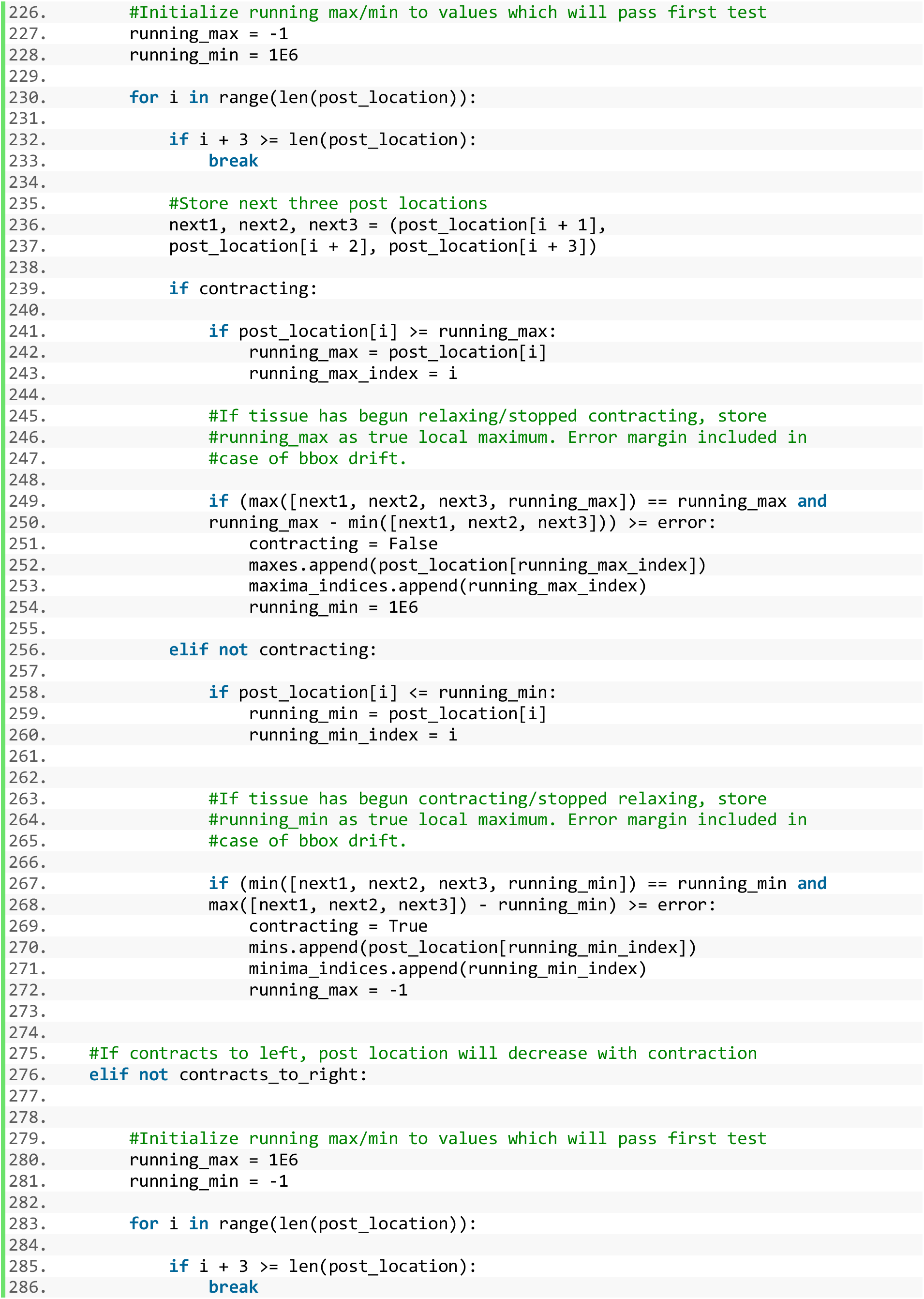
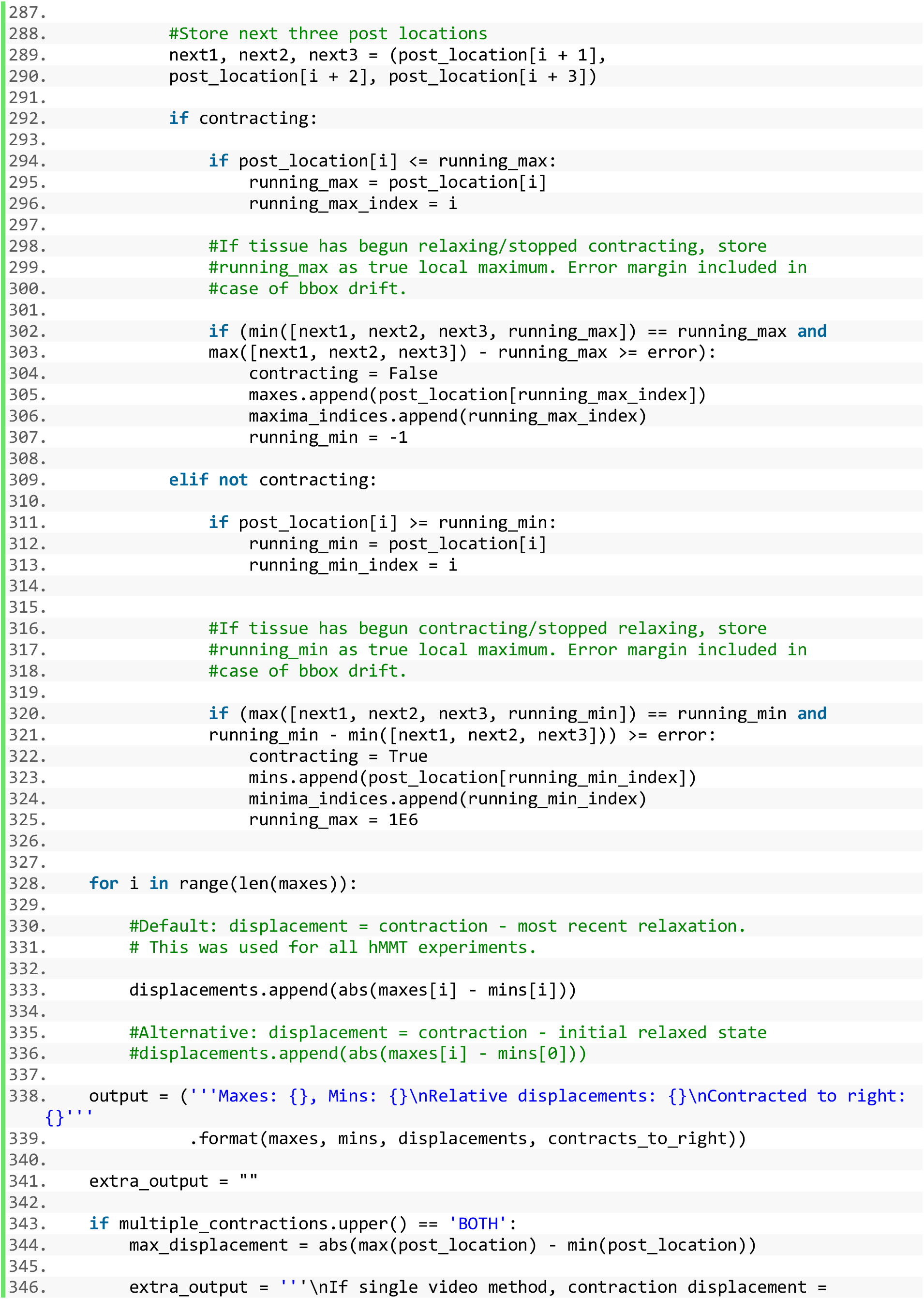
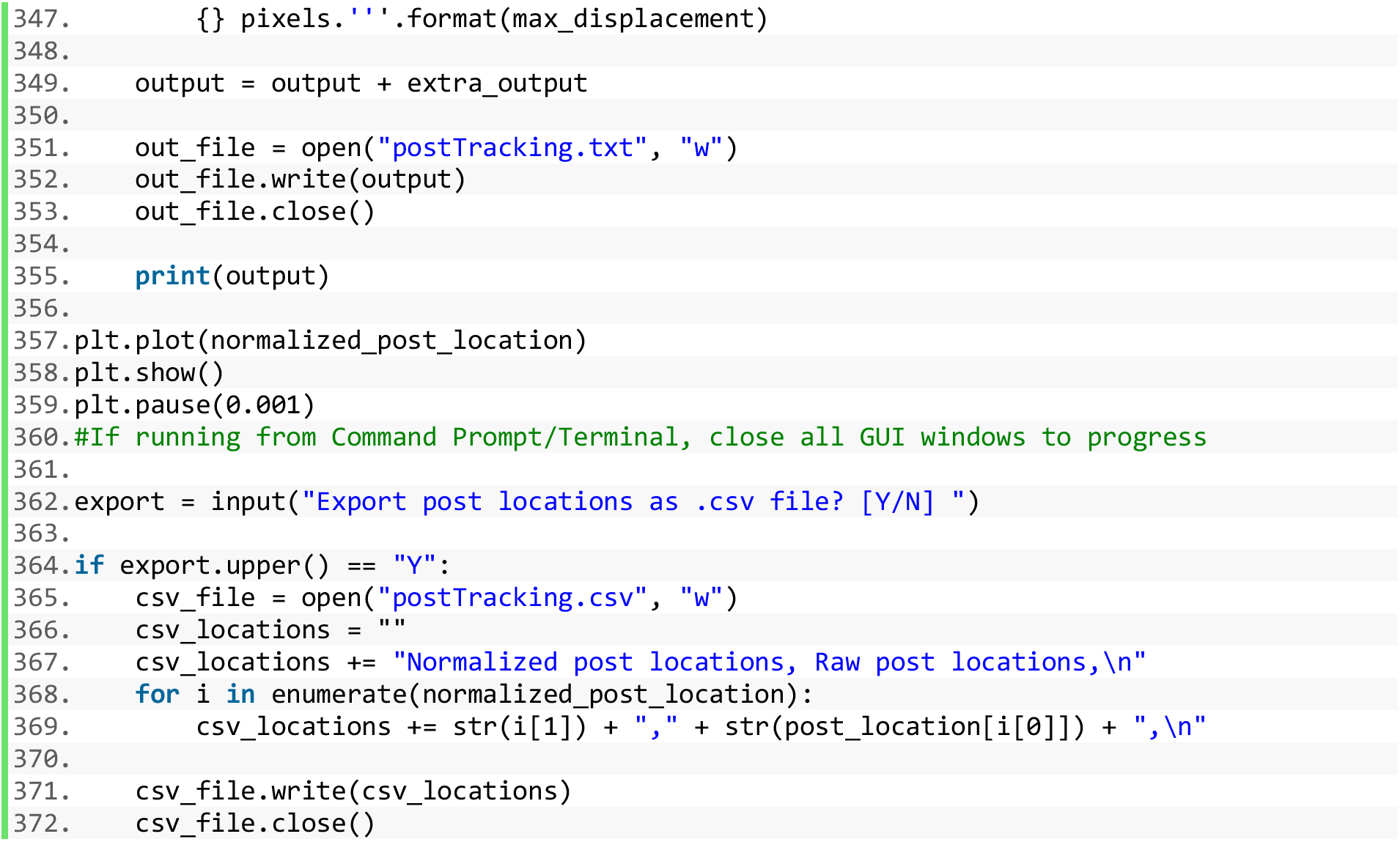

## REFERENCES

Agrawal, G., Aung, A., and Varghese, S. (2017). Skeletal muscle-on-a-chip: an in vitro model to evaluate tissue formation and injury. Lab. Chip 17, 3447–3461.

Alnaqeeb, M.A., Al Zaid, N.S., and Goldspink, G. (1984). Connective tissue changes and physical properties of developing and ageing skeletal muscle. J. Anat. 139 (Pt 4), 677–689.

Argilés, J.M., Busquets, S., Stemmler, B., and López-Soriano, F.J. (2014). Cancer cachexia: understanding the molecular basis. Nat. Rev. Cancer 14, 754–762.

Bakooshli, M.A., Lippmann, E.S., Mulcahy, B., Iyer, N., Nguyen, C.T., Tung, K., Stewart, B.A., Dorpel, H. van den Fuehrmann, T., Shoichet, M.S., et al. (2018). A three-dimensional culture model of innervated human skeletal muscle enables studies of the adult neuromuscular junction and disease modeling. BioRxiv 275545.

Ballantyne, J.C., and Chang, Y.C. (1997). The impact of choice of muscle relaxant on postoperative recovery time: A retrospective study. Anesth. Analg.

Barreto, R., Waning, D.L., Gao, H., Liu, Y., Zimmers, T.A., and Bonetto, A. (2016). Chemotherapy-related cachexia is associated with mitochondrial depletion and the activation of ERK1/2 and p38 MAPKs. Oncotarget 7, 43442–43460.

Blau, H.M., and Webster, C. (1981). Isolation and characterization of human muscle cells. Proc. Natl. Acad. Sci. 78, 5623–5627.

Bodine, S.C., Stitt, T.N., Gonzalez, M., Kline, W.O., Stover, G.L., Bauerlein, R., Zlotchenko, E., Scrimgeour, A., Lawrence, J.C., Glass, D.J., et al. (2001). Akt/mTOR pathway is a crucial regulator of skeletal muscle hypertrophy and can prevent muscle atrophy in vivo. Nat. Cell Biol. 3, 1014–1019.

Bowman, W.C. (2006). Neuromuscular block. Br. J. Pharmacol.

Chabot, G.G. (1997). Clinical pharmacokinetics of irinotecan. Clin. Pharmacokinet. 33, 245–259.

Chal, J., Al Tanoury, Z., Hestin, M., Gobert, B., Aivio, S., Hick, A., Cherrier, T., Nesmith, A.P., Parker, K.K., and Pourquié, O. (2016). Generation of human muscle fibers and satellite-like cells from human pluripotent stem cells in vitro. Nat. Protoc. 11, 1833–1850.

Chen, T.-W., Wardill, T.J., Sun, Y., Pulver, S.R., Renninger, S.L., Baohan, A., Schreiter, E.R., Kerr, R.A., Orger, M.B., Jayaraman, V., et al. (2013). Ultrasensitive fluorescent proteins for imaging neuronal activity. Nature 499, 295–300.

Coleman, M.E., DeMayo, F., Yin, K.C., Lee, H.M., Geske, R., Montgomery, C., and Schwartz, R.J. (1995). Myogenic vector expression of insulin-like growth factor I stimulates muscle cell differentiation and myofiber hypertrophy in transgenic mice. J. Biol. Chem. 270, 12109–12116.

Conroy, T., Desseigne, F., Ychou, M., Bouché, O., Guimbaud, R., Bécouarn, Y., Adenis, A., Raoul, J.-L., Gourgou-Bourgade, S., de la Fouchardière, C., et al. (2011). FOLFIRINOX versus Gemcitabine for Metastatic Pancreatic Cancer. N. Engl. J. Med. 364, 1817–1825.

DiMasi, J.A., Hansen, R.W., and Grabowski, H.G. (2003). The price of innovation: new estimates of drug development costs. J. Health Econ. 22, 151–185.

Dimitriu, C., Martignoni, M.E., Bachmann, J., Fröhlich, B., Tintărescu, G., Buliga, T., Lică, I., Constantinescu, G., Beuran, M., and Friess, H. (2005). Clinical impact of cachexia on survival and outcome of cancer patients. Romanian J. Intern. Med. Rev. Roum. Med. Interne 43, 173–185.

Dodson, S., Baracos, V.E., Jatoi, A., Evans, W.J., Cella, D., Dalton, J.T., and Steiner, M.S. (2011). Muscle Wasting in Cancer Cachexia: Clinical Implications, Diagnosis, and Emerging Treatment Strategies. Annu. Rev. Med. 62, 265–279.

Duance, V.C., Stephens, H.R., Dunn, M., Bailey, A.J., and Dubowitz, V. (1980). A role for collagen in the pathogenesis of muscular dystrophy? Nature 284, 470–472.

Eberli, D., Soker, S., Atala, A., and Yoo, J.J. (2009). Optimization of human skeletal muscle precursor cell culture and myofiber formation in vitro. Methods 47, 98–103.

Gaschen, F.P., Hoffman, E.P., Gorospe, J.R., Uhl, E.W., Senior, D.F., Cardinet, G.H., and Pearce, L.K. (1992). Dystrophin deficiency causes lethal muscle hypertrophy in cats. J. Neurol. Sci. 110, 149–159.

Hinds, S., Bian, W., Dennis, R.G., and Bursac, N. (2011). The Role of Extracellular Matrix Composition in Structure and Function of Bioengineered Skeletal Muscle. Biomaterials 32, 3575–3583.

Jacquemin, V., Furling, D., Bigot, A., Butler-Browne, G.S., and Mouly, V. (2004). IGF-1 induces human myotube hypertrophy by increasing cell recruitment. Exp. Cell Res. 299, 148–158.

Juhas, M., Engelmayr, G.C., Fontanella, A.N., Palmer, G.M., and Bursac, N. (2014). Biomimetic engineered muscle with capacity for vascular integration and functional maturation in vivo. Proc. Natl. Acad. Sci. 201402723.

Keil, A., Frese-Schaper, M., Steiner, S.K., Körner, M., Schmid, R.A., and Frese, S. (2015). The topoisomerase I inhibitor irinotecan and the tyrosyl-DNA phosphodiesterase 1 inhibitor furamidine synergistically suppress murine lupus nephritis. Arthritis Rheumatol. 67, 1858–1867.

Lauretani, F., Russo, C.R., Bandinelli, S., Bartali, B., Cavazzini, C., Di Iorio, A., Corsi, A.M., Rantanen, T., Guralnik, J.M., and Ferrucci, L. (2003). Age-associated changes in skeletal muscles and their effect on mobility: an operational diagnosis of sarcopenia. J. Appl. Physiol. Bethesda Md 1985 95, 1851–1860.

Lee, P.H.U., and Vandenburgh, H.H. (2013). Skeletal Muscle Atrophy in Bioengineered Skeletal Muscle: A New Model System. Tissue Eng. Part A 19, 2147–2155.

Madden, L., Juhas, M., Kraus, W.E., Truskey, G.A., and Bursac, N. (2015). Bioengineered human myobundles mimic clinical responses of skeletal muscle to drugs. ELife 4, e04885.

Maffioletti, S.M., Sarcar, S., Henderson, A.B.H., Mannhardt, I., Pinton, L., Moyle, L.A., Steele-Stallard, H., Cappellari, O., Wells, K.E., Ferrari, G., et al. (2018). Three-Dimensional Human iPSC-Derived Artificial Skeletal Muscles Model Muscular Dystrophies and Enable Multilineage Tissue Engineering. Cell Rep. 23, 899–908.

Maltzahn, J. von Renaud, J.-M., Parise, G., and Rudnicki, M.A. (2012). Wnt7a treatment ameliorates muscular dystrophy. Proc. Natl. Acad. Sci. 109, 20614–20619.

McGreevy, J.W., Hakim, C.H., McIntosh, M.A., and Duan, D. (2015). Animal models of Duchenne muscular dystrophy: from basic mechanisms to gene therapy. Dis. Model. Mech. 8, 195–213.

Moysan, E., Bastiat, G., and Benoit, J.P. (2013). Gemcitabine versus modified gemcitabine: A review of several promising chemical modifications. Mol. Pharm. 10, 430–444.

Musarò, A., McCullagh, K., Paul, A., Houghton, L., Dobrowolny, G., Molinaro, M., Barton, E.R., Sweeney, H.L., and Rosenthal, N. (2001). Localized Igf-1 transgene expression sustains hypertrophy and regeneration in senescent skeletal muscle. Nat. Genet. 27, 195–200.

Osaki, T., Uzel, S.G.M., and Kamm, R.D. (2018). Microphysiological 3D model of amyotrophic lateral sclerosis (ALS) from human iPS-derived muscle cells and optogenetic motor neurons. Sci. Adv. 4.

Periasamy, M., Maurya, S.K., Sahoo, S.K., Singh, S., Sahoo, S.K., Reis, F.C.G., and Bal, N.C. (2017). Role of SERCA Pump in Muscle Thermogenesis and Metabolism. Compr. Physiol. 7, 879–890.

Powell, C.A., Smiley, B.L., Mills, J., and Vandenburgh, H.H. (2002). Mechanical stimulation improves tissue-engineered human skeletal muscle. Am. J. Physiol. - Cell Physiol. 283, C1557–C1565.

Rao, L., Qian, Y., Khodabukus, A., Ribar, T., and Bursac, N. (2018). Engineering human pluripotent stem cells into a functional skeletal muscle tissue. Nat. Commun. 9, 126.

Rommel, C., Clarke, B.A., Zimmermann, S., Nuñez, L., Rossman, R., Reid, K., Moelling, K., Yancopoulos, G.D., and Glass, D.J. (1999). Differentiation Stage-Specific Inhibition of the Raf-MEK-ERK Pathway by Akt. Science 286, 1738–1741.

Rommel, C., Bodine, S.C., Clarke, B.A., Rossman, R., Nunez, L., Stitt, T.N., Yancopoulos, G.D., and Glass, D.J. (2001). Mediation of IGF-1-induced skeletal myotube hypertrophy by PI(3)K/Akt/mTOR and PI(3)K/Akt/GSK3 pathways. Nat. Cell Biol. 3, 1009–1013.

Rybalka, E., Timpani, C.A., Cheregi, B.D., Sorensen, J.C., Nurgali, K., and Hayes, A. (2018). Chemotherapeutic agents induce mitochondrial superoxide production and toxicity but do not alter respiration in skeletal muscle in vitro. Mitochondrion 42, 33–49.

Sakar, M.S., Neal, D., Boudou, T., Borochin, M.A., Li, Y., Weiss, R., Kamm, R.D., Chen, C.S., and Asada, H.H. (2012). Formation and optogenetic control of engineered 3D skeletal muscle bioactuators. Lab. Chip 12, 4976–4985.

Semsarian, C., Sutrave, P., Richmond, D.R., and Graham, R.M. (1999). Insulin-like growth factor (IGF-I) induces myotube hypertrophy associated with an increase in anaerobic glycolysis in a clonal skeletal-muscle cell model. Biochem. J. 339 (Pt 2), 443–451.

Shao, G., Wu, J., Cai, Z., and Wang, W. (2012). Fabrication of elastomeric high-aspect-ratio microstructures using polydimethylsiloxane (PDMS) double casting technique. Sens. Actuators Phys. 178, 230–236.

Smith, A.S.T., Long, C.J., Pirozzi, K., Najjar, S., McAleer, C., Vandenburgh, H.H., and Hickman, J.J. (2014). A multiplexed chip-based assay system for investigating the functional development of human skeletal myotubes in vitro. J. Biotechnol. 185, 15–18.

Staffa, J.A., Chang, J., and Green, L. (2002). Cerivastatin and Reports of Fatal Rhabdomyolysis. N. Engl. J. Med. 346, 539–540.

Stevenson, E.J., Koncarevic, A., Giresi, P.G., Jackman, R.W., and Kandarian, S.C. (2005). Transcriptional profile of a myotube starvation model of atrophy. J. Appl. Physiol. Bethesda Md 1985 98, 1396–1406.

Sun, L., Quan, X.-Q., and Yu, S. (2015). An Epidemiological Survey of Cachexia in Advanced Cancer Patients and Analysis on Its Diagnostic and Treatment Status. Nutr. Cancer 67, 1056–1062.

Tamraz, B., Fukushima, H., Wolfe, A.R., Kaspera, R., Totah, R.A., Floyd, J.S., Ma, B., Chu, C., Marciante, K.D., Heckbert, S.R., et al. (2013). OATP1B1-related drug-drug and drug-gene interactions as potential risk factors for cerivastatin-induced rhabdomyolysis. Pharmacogenet. Genomics 23, 355–364.

Thomason, D.B., and Booth, F.W. (1990). Atrophy of the soleus muscle by hindlimb unweighting. J. Appl. Physiol. Bethesda Md 1985 68, 1–12.

Vandenburgh, H., Shansky, J., Benesch-Lee, F., Barbata, V., Reid, J., Thorrez, L., Valentini, R., and Crawford, G. (2008). Drug-screening platform based on the contractility of tissue-engineered muscle. Muscle Nerve 37, 438–447.

Vandenburgh, H., Shansky, J., Benesch-Lee, F., Skelly, K., Spinazzola, J.M., Saponjian, Y., and Tseng, B.S. (2009). Automated drug screening with contractile muscle tissue engineered from dystrophic myoblasts. FASEB J. 23, 3325–3334.

Vandenburgh, H.H., Karlisch, P., Shansky, J., and Feldstein, R. (1991). Insulin and IGF-I induce pronounced hypertrophy of skeletal myofibers in tissue culture. Am. J. Physiol. 260, C475–484.

Velloso, C.P. (2008). Regulation of muscle mass by growth hormone and IGF-I. Br. J. Pharmacol. 154, 557–568.

Viguerie, N., Picard, F., Hul, G., Roussel, B., Barbe, P., Iacovoni, J.S., Valle, C., Langin, D., and Saris, W.H.M. (2012). Multiple effects of a short-term dexamethasone treatment in human skeletal muscle and adipose tissue. Physiol. Genomics 44, 141–151.

Ye, F., Mathur, S., Liu, M., Borst, S.E., Walter, G.A., Sweeney, H.L., and Vandenborne, K. (2013). Overexpression of insulin-like growth factor-1 attenuates skeletal muscle damage and accelerates muscle regeneration and functional recovery after disuse. Exp. Physiol. 98, 1038–1052.

Young, J., Margaron, Y., Fernandes, M., Duchemin-Pelletier, E., Michaud, J., Flaender, M., Lorintiu, O., Degot, S., and Poydenot, P. (2018). MyoScreen, a High-Throughput Phenotypic Screening Platform Enabling Muscle Drug Discovery. SLAS Discov. Adv. Life Sci. RD 23, 790–806.

